# Live-cell 3D single-molecule tracking reveals how NuRD modulates enhancer dynamics

**DOI:** 10.1101/2020.04.03.003178

**Authors:** S Basu, O Shukron, D Hall, P Parutto, A Ponjavic, D Shah, W Boucher, D Lando, W Zhang, N Reynolds, LH Sober, A Jartseva, R Ragheb, X Ma, J Cramard, R Floyd, G Brown, J Balmer, TA Drury, AR Carr, L-M Needham, A Aubert, G Communie, K Gor, M Steindel, L Morey, E Blanco, T Bartke, L Di Croce, I Berger, C Schaffitzel, SF Lee, TJ Stevens, D Klenerman, BD Hendrich, D Holcman, ED Laue

**Affiliations:** Department of Biochemistry, University of Cambridge, 80 Tennis Court Road, Cambridge CB2 1GA, United Kingdom; Wellcome-MRC Cambridge Stem Cell Institute, Jeffrey Cheah Biomedical Centre, Puddicombe Way, Cambridge CB2 0AW, United Kingdom; Department of Applied Mathematics and Computational Biology, Ecole Normale Supérieure, 75005 Paris, France; Yusuf Hamied Department of Chemistry, University of Cambridge, Lensfield Road, Cambridge, CB2 1EW, United Kingdom; The European Molecular Biology Laboratory EMBL, 71 Avenue des Martyrs, 38000 Grenoble, France; Helmholtz Zentrum München, German Research Center for Environmental Health, Institute of Functional Epigenetics, Ingolstädter Landstr. 1, 85764 Neuherberg, Germany; Centre for Genomic Regulation (CRG), The Barcelona Institute of Science and Technology, Dr. Aiguader 88, Barcelona 08003, Spain; Institució Catalana de Recerca i Estudis Avançats (ICREA), Pg. Lluis Companys 23, Barcelona, Spain; School of Biochemistry, University of Bristol, 1 Tankard’s Close, Bristol BS8 1TD, United Kingdom; MRC Laboratory of Molecular Biology, Francis Crick Avenue, Cambridge Biomedical Campus, Cambridge CB2 0QH

## Abstract

Enhancer-promoter dynamics are crucial for the spatiotemporal control of gene expression, but it remains unclear whether these dynamics are controlled by chromatin regulators, such as the nucleosome remodelling and deacetylase (NuRD) complex. The NuRD complex binds to all active enhancers to modulate transcription and here we use Hi-C experiments to show that it blurs TAD boundaries and increases the proximity of intermediate-range (~1 Mb) genomic sequences and enhancer-promoter interactions. To understand whether NuRD alters the dynamics of 3D genome structure, we developed an approach to segment and extract key biophysical parameters from 3D single-molecule trajectories of the NuRD complex determined using live-cell imaging. Unexpectedly, this revealed that the intact NuRD complex decompacts chromatin structure and makes NuRD-bound enhancers move faster, increasing the overall volume of the nucleus that these key regulatory regions explore. Interestingly, we also uncovered a rare fast-diffusing state of NuRD bound enhancers that exhibits directed motion. The NuRD complex reduces the amount of time that enhancers remain in this fast-diffusing state, which we propose might otherwise re-organise enhancer-promoter proximity. Thus, we uncover an intimate connection between a chromatin remodeller and the spatial dynamics of the local regions of the genome to which it binds.

## Introduction

3D genome organisation and chromatin dynamics are thought to be crucial for the spatiotemporal control of gene expression. However, little is known about the multi-scale chromatin dynamics of cis-regulatory elements or how they relate to genome organisation. To probe the dynamics and organisation of these elements at a single-cell level two complementary methods are typically used – either live-cell imaging^1–5^ or single nucleus versions of chromosome conformation capture experiments^6–13^ (such as Hi-C), which reveal snapshots of the structure of a dynamic 3D genome in different individual fixed cells^6–17^. Whether chromatin regulators modulate these dynamics or how these dynamics relate to genome structure remains unclear.

The nucleosome remodelling and deacetylase (NuRD) complex is a highly conserved 1 MDa multi-subunit protein complex which binds to all active enhancers^18^. NuRD combines two key enzymatic activities – nucleosome remodelling *via* its helicase-containing ATPase (predominantly CHD4 in embryonic stem (ES) cells and lysine deacetylation *via* its HDAC1/2 subunits^17–23^: these activities are thought to be present in two sub-complexes. HDAC1/2 associate, along with the histone chaperones RBBP4/7, with the core scaffold proteins MTA1/2/3 to form a stable sub-complex with deacetylase activity^24^. The nucleosome remodeller CHD4 interacts with chromatin by itself and also forms a second sub-complex with GATAD2A/B and DOC1 (CDK2AP1)^24–26^. The methyl-CpG DNA binding domain proteins MBD2/3 interact directly with both the deacetylase sub-complex and GATAD2A/B^24,27–29^, and thus play a critical role in linking the CHD4 remodeller and HDAC sub-complexes together to assemble the intact holo-NuRD complex (**Figure 1a**).

**Figure 1.**
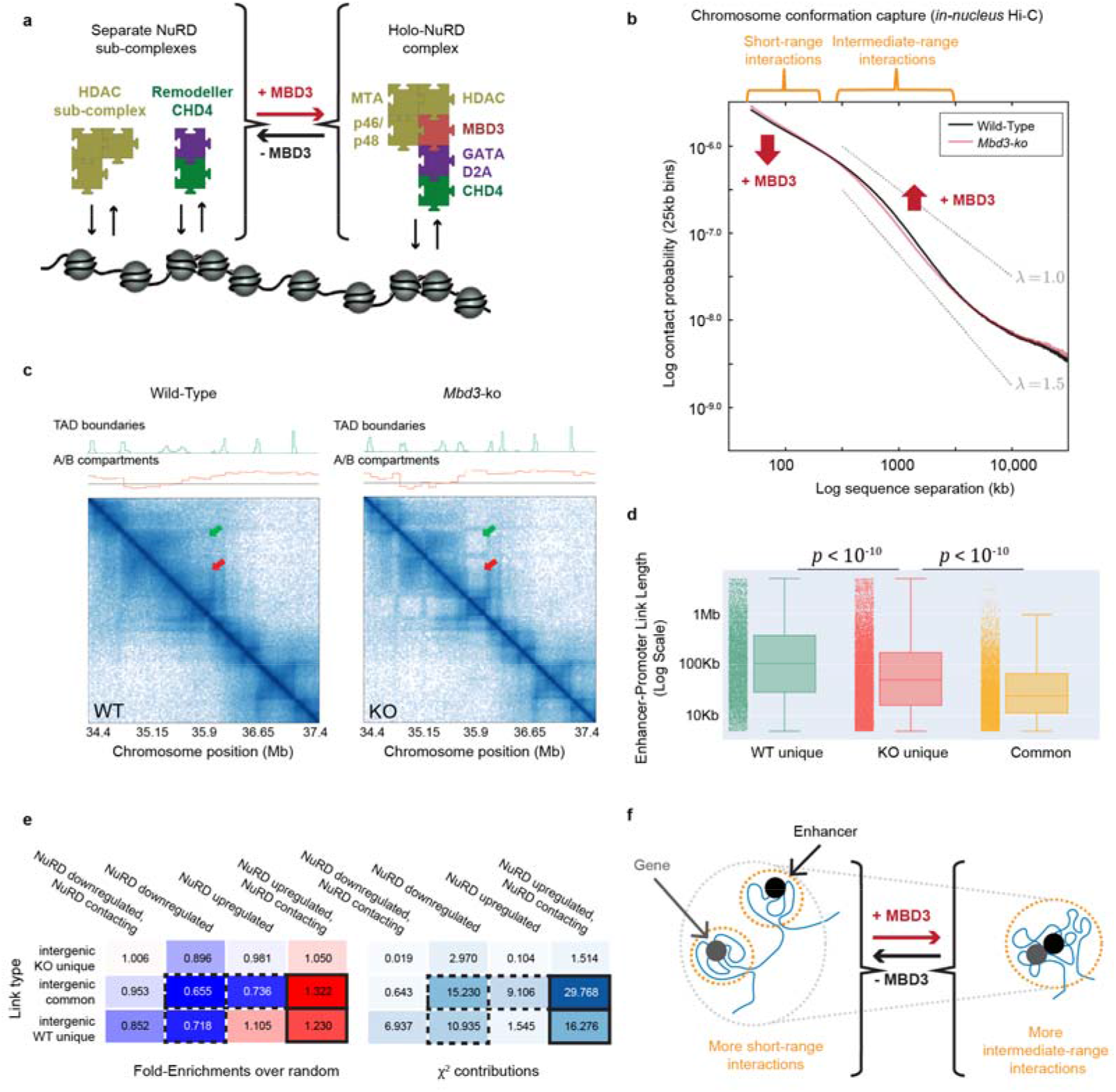
MBD3-dependent assembly of the NuRD complex increases Mb-range genome interactions. (a) Schematic representation of the NuRD complex interacting with chromatin in the presence and absence of MBD3. (b) Log-log plots of contact probability as a function of genomic sequence separation (averaged across the genome), derived from *in-nucleus* Hi-C experiments of *Mbd3*-ko (red) and wild-type (black) ES cells, shows a significant (30%) increase in intermediate-range (~1 Mb) contacts in wild-type cells (p<10^−17^, Wilcoxon rank sum test, see **Extended Data Figure 1b**). (c) Representative plots showing part of the Hi-C contact map for Chromosome 1 – the density of contacts is indicated by the colour intensity. Some TAD boundaries are weakened resulting in an increased density of contacts between adjacent TADs, both within and between A/B compartments (red and green arrows) – see also **Extended Data Figures 1g-i** for genome-wide comparisons. (d) Boxplots showing intra-chromosomal enhancer-promoter link lengths, determined using a modified version of the activity-by-contact algorithm^50^, present in both *Mbd3*-ko and wild-type ES cells (orange), in only *Mbd3*-ko cells (red) or in only wild-type cells (green). The bars on the left of each plot show all the data. (e) (Left) Fold-enrichment and (Right) chi-squared test when intergenic enhancer-promoter interactions found in both wild-type and *Mbd3*-ko cells, as well as those found uniquely in either the *Mbd3-*ko or the wild-type, are correlated with genes that are up−/down-regulated in the presence of intact NuRD. Interactions where NuRD bound enhancers can be seen to contact the promoter, that are found in both the *Mbd3-*ko and the wild-type, or uniquely in the wild-type, are enriched at up-regulated genes. In contrast, there is a depletion in interactions between intergenic enhancers and the promoter at down-regulated genes. Enriched and depleted interactions are coloured red and blue, respectively, and highlighted using solid and dashed black boxes – see also **Extended Data Figure 3c** for an example of changes in enhancer-promoter contacts. (f) Schematic interpretation of the results of the Hi-C experiments, showing that in the *Mbd3* knockout there are more short range and fewer intermediate range chromatin interactions.

Assembly of the intact holo-NuRD complex is critical for controlling cell fate transitions. Knockout of *Mbd3*, which disrupts intact NuRD complex assembly, leads to only moderate up- or down-regulation (‘fine-tuning’) of transcription levels, but this modulation prevents ES cell lineage commitment^30–33^. Nucleosome remodelling by NuRD regulates transcription factor and RNA polymerase II binding at active enhancers^18^, but whether this impacts enhancer dynamics or enhancer-promoter interactions has remained unclear. Here, to understand whether enhancer dynamics are regulated by this crucial chromatin remodeller, we combine Hi-C and live-cell single-molecule tracking. Specifically, we exploit the ability to unlink the chromatin remodelling and deacetylase subunits of intact NuRD by deleting *Mbd3* to explore the function of the intact NuRD complex.

### NuRD complex assembly increases contacts between intermediate-range genomic sequences

We initially set out to understand whether assembly of the intact NuRD complex, mediated by MBD3, might affect genome architecture by carrying out chromosome conformation capture (*in-nucleus* Hi-C) experiments on wild-type and previously characterised^31^ *Mbd3* knockout (*Mbd3-ko)* ES cells. This strategy, which exploits the fact that MBD2 (and thus the MBD2-linked holo-NuRD complex) is expressed at low levels in ES cells and cannot rescue the *Mbd3* deletion^34,35^, allowed us to specifically perturb NuRD complex function. We obtained high-quality Hi-C contact maps for both wild-type and *Mbd3-ko* ES cells after combining our wild-type data with previously published^36^ (and consistent) datasets (**Extended Data Figure 1a**). As previously observed^6,8–17,37^, these contact maps showed that the genome is segregated into: *1)* A and B compartments (broadly regions containing more or fewer genes respectively); *2)* megabase-scale topologically associating domains (TADs), which have a higher frequency of intra-domain chromatin interactions; and *3)* loops where specific genomic regions contact each other, mediated e.g. *via* CTCF/Cohesin binding.

Comparison of the *Mbd3-ko* and wild-type Hi-C data showed that assembly of the intact NuRD complex leads to a ~30 % genome-wide increase in the probability of intermediate-range contacts at the scale of TADs (500 kb to 3 Mb) (p < 10^−17^, **Figure 1b**, **Extended Data Figure 1b**). We found that ~77% and 17% of NuRD-regulated genes^38^ remain in the A and B compartments, respectively, with only a small proportion (~3%) moving from the A to the B compartment or vice versa. The genome-wide increase in mean contact length/region binned (**Extended Data Figure 1c**) was most noticeable for regions containing NuRD-regulated genes that are within the A compartment in both *Mbd3-ko* and wild-type cells (KO-A, WT-A; see **Extended Data Figure 1d**). We also found that promoters that are down-regulated in the presence of intact NuRD are enriched (chi-squared p-value: < 1e-10, odds-ratio: 1.16) amongst genes that remain in the A compartment and significantly depleted (chi-squared p-value: < 1e-10, odds-ratio: 0.43) amongst those that remain within the B compartment. Further classification showed that promoters displaying significant MBD3 (i.e. NuRD) ChIP-seq signal [which our unpublished experiments (RR, NR & BDH) suggest represents association with a NuRD-bound enhancer] are also significantly enriched amongst genes that remain in the A compartment (p < 10^−10^, odds-ratio: 1.27) and significantly depleted amongst those that remain within the B compartment (p < 10^−10^, odds-ratio: 0.41) (**Extended Data Figures 1e,f**). Thus, NuRD-bound enhancers tend to contact promoters to down-regulate genes in the A (euchromatic) compartment.

Comparison of the Hi-C contact maps for *Mbd3-ko* and wild-type cells showed that some boundaries between TAD’s and A/B compartments are weaker in cells containing the intact NuRD complex (**Figure 1c** and **Extended Data Figure 1g**), with promoters that contact NuRD-bound enhancers having an increased number of cross-compartment contacts (**Extended Data Figure 1h**) thereby decreasing TAD insulation (**Extended Data Figure 1i**). Although we could not detect a significant correlation between the A/B compartmentalisation of a gene and levels of transcription (**Extended Data Figure 1j**), these results show that NuRD-regulated genes tend to be at A/B compartment boundaries^39,40^. The blurring of TAD and A/B compartment boundaries suggested that NuRD might alter CTCF/Cohesin binding^40,41^. We therefore carried out Cut&Run experiments to study chromatin binding of CTCF and SMC3 (a subunit of the Cohesin complex) – which play a key role in loop and TAD formation^13,42–47^. On assembly of the intact NuRD complex we found there was a redistribution of both CTCF and SMC3 binding – the effect on SMC3 being more pronounced – with both CTCF and SMC3 peaks appearing and disappearing (**Extended Data Figure 2b**; see also **Extended Data Figure 2e** for an example gene). We found that NuRD-regulated genes are more likely to be found near to CTCF/Cohesin binding sites that form when the intact NuRD complex assembles – this was more noticeable for genes that were upregulated in wild-type vs *Mbd3-ko* cells and whose promoters contact NuRD bound enhancers (**Extended Data Figure 2d**). Interestingly, we also found that genes that are regulated in a similar way by NuRD (i.e. up or down) tend to cluster with each other – see for example the *Htra1* gene, which is found in a region with other significantly downregulated genes (**Extended Data Figure 2e**).

Because changes in TADs and CTCF-Cohesin loops are thought to influence enhancer-promoter proximity^45,48,49^, we used a modified version of the activity-by-contact algorithm^50^ to determine changes in enhancer-promoter interactions in *Mbd3-ko* vs wild-type cells (**Extended Data Figures 3a,b**). Using active enhancers and promoters defined using H3K27ac and H3K4me3 ChIP-seq profiles^18^ (see Online Methods), this analysis revealed that enhancer-promoter interactions that only occur in the presence of the intact NuRD complex tend to link together genomic regions separated by longer distances than those that occur in the absence of MBD3 (**Figure 1d**). In addition, when we considered enhancer-promoter interactions and transcription levels in wild-type compared to *Mbd3-ko* cells^38^, we found more contacts between intergenic NuRD bound enhancers and promoters that are upregulated by NuRD, and fewer contacts between intergenic enhancers and promoters that are downregulated by NuRD (**Figure 1e**).

### A novel trajectory segmentation algorithm for extracting NuRD complex dynamics and studying chromatin movement

To provide a mechanistic understanding of how assembly of the intact NuRD complex might increase mixing of intermediate-range genome sequences, we carried out live-cell 3D single-molecule tracking of NuRD complex subunits in the presence and absence of MBD3. We generated knock-in ES cell lines expressing the endogenous *Chd4*, *Mbd3* and *Mta2* genes fused with C-terminal HaloTags and confirmed that the tags did not prevent NuRD complex assembly, although subtle changes in expression were observed (**Extended Data Figure 4**). We used a double-helix point spread function microscope^25^ to generate 3D tracks of single NuRD-HaloTag-JF_549_ complexes through a 4 μm slice of the nucleus, so that we could reliably determine biophysical parameters (**Figure 2a**). (Tracking in 2D reduces the length of trajectories because molecules can move in/out of the imaging plane during the experiment.) We recorded trajectories at two distinct temporal regimes, 20 ms and 500 ms. Recording at 20 ms time resolution allows the detection of both freely diffusing and chromatin bound proteins^25^, and can thus be used to extract the chromatin binding kinetics of NuRD complexes (**Figure 2b**). In contrast, at a 500 ms time resolution, ‘motion blurring’ substantially reduces the detection of freely diffusing molecules, allowing us to focus on the slower sub-diffusive chromatin bound NuRD complexes^1^ (**Figure 2c**).

**Figure 2.**
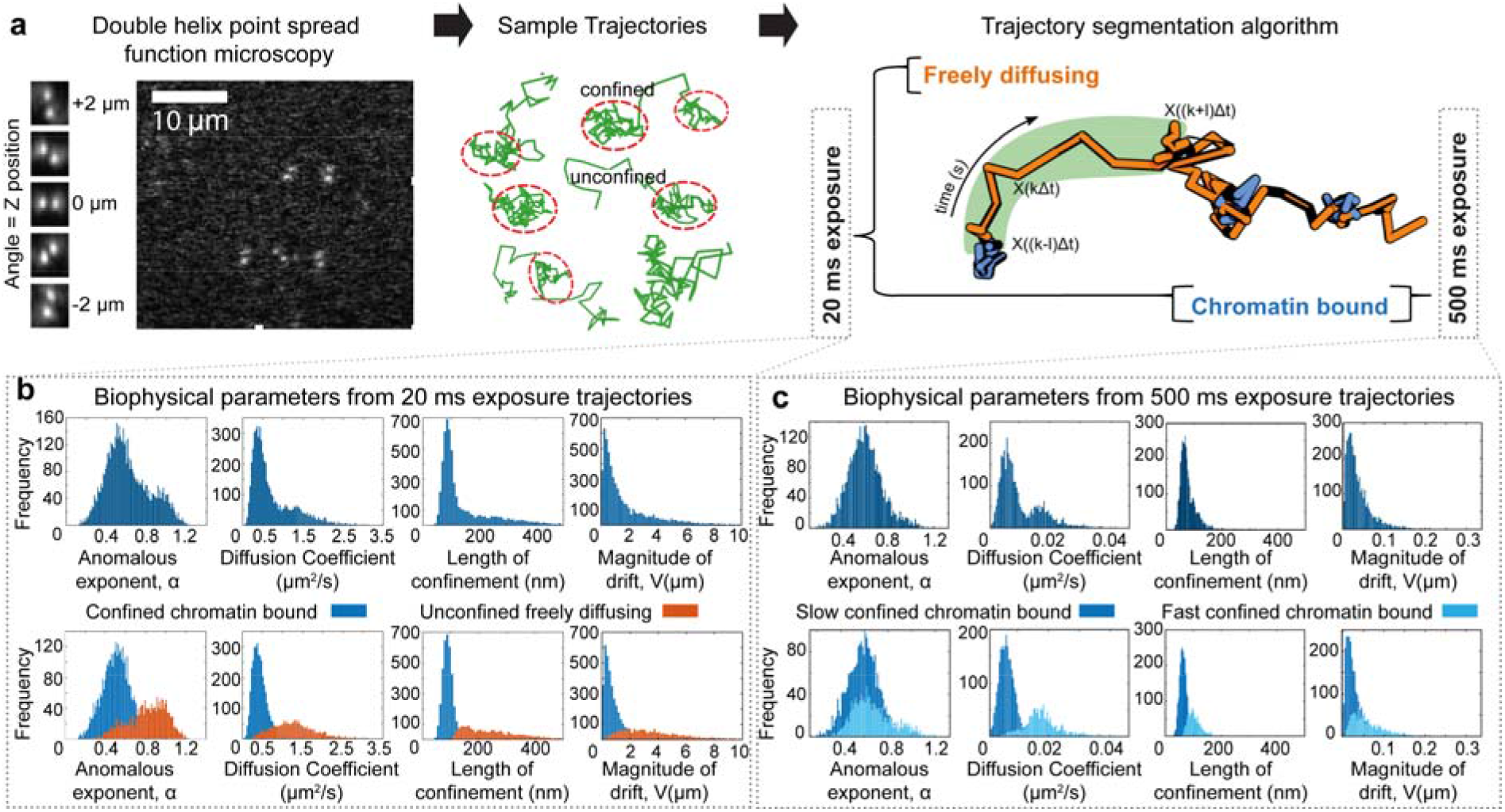
Live cell imaging to study NuRD complex binding kinetics and function. (a) (Left) Single JF_549_-HaloTagged CHD4 molecules in the NuRD complex are tracked in 3D using a double-helix point spread function microscope – two puncta are recorded for each fluorophore with their midpoint providing the lateral x, y position and the angle between them representing the axial position in z relative to the nominal focal plane. (Middle) Examples of extracted single particle trajectories from 20 ms exposure imaging show periods of free unconfined and confined diffusion. (Right) Example trajectory of a single CHD4 molecule from 20 ms exposure imaging showing a sliding window of 11 frames (shaded green) from which the biophysical parameters are calculated. The resulting segmentation, into confined chromatin bound (blue) and unconfined freely diffusing (orange) sub-trajectories, has been used to colour the trajectory. 500 ms exposure imaging, on the other hand, can be used to extract slow and fast sub-diffusive motion of chromatin bound CHD4 molecules. (b, c) Distribution of the four biophysical parameters extracted from sliding windows within the single-molecule trajectories of CHD4 taken with either (b) 20 ms or (c) 500 ms exposures – (top) before and (bottom) after classification based on the anomalous exponent α, the apparent diffusion coefficient D, the length of confinement Lc, and the drift magnitude, norm∥V∥ of the mean velocity. Each dataset was produced by imaging 30 cells, and the parameters were extracted from 5,557 and 15,528 trajectories, respectively (see **Extended Data Figure 5** and the Online Methods for more detail).

To extract dynamic biophysical parameters, we developed a machine learning method (a Gaussian mixture model) to segment the single molecule trajectories into different classes by studying their behaviour over a sliding window of 11 consecutive images (see **Figure 2** and **Extended Data Figure 5**). To understand how NuRD affects chromatin structure, we estimated from each sub-trajectory not just the apparent diffusion coefficient (as previously used for classifying sub-trajectories^51,52^) but also the anomalous exponent α, the localisation length Lc, and the drift magnitude V^53^. The anomalous exponent α, defined as [mean squared displacement ∝ *t*^α^], is particularly informative. Diffusing proteins are characterised by an anomalous exponent close to 1 whereas chromatin bound proteins exhibit a lower anomalous exponent^3,53,54^, whose value is representative of the condensation state of the chromatin^53,55^. In addition, values of α that are higher than 1 can represent energy-dependent directed motion. The localisation length Lc of chromatin bound proteins is also informative as it reflects the spatial scale that the molecules explore within the nucleus. The localisation length Lc is dependent on a range of parameters such as nucleosome density and CTCF/Cohesin-mediated looping interactions: for example, Lc is larger at lower nucleosome densities when the linker length between nucleosomes is longer^55,56^. Finally, by computing the magnitude *V_i_* (k) of the drift vector *V_i_* in three dimensions we can characterise directed movement of a molecule. (Further details of the approach – and of the simulations we carried out to test the algorithm – are described in the Online Methods.)

Analysis of the 20 ms exposure trajectories of single CHD4 molecules using our approach revealed two diffusion states (**Figure 2b**). We identified a fast unconfined state that was freely diffusing with an α of 0.94 ± 0.12 and an apparent diffusion coefficient of 1.3 ± 0.3 μm^2^/s (matching previous observations^24,25^), as well as a confined chromatin bound state characterised by sub-diffusive motion with an α of 0.51 ± 0.02 and an apparent diffusion coefficient of 0.43 ± 0.03 μm^2^/s. Similar results were obtained when segmenting the trajectories of two other NuRD complex components, MBD3 and MTA2 (**Extended Data Figure 6c**). To demonstrate that our approach can reliably determine relative differences in anomalous exponent for freely diffusing and sub-diffusive chromatin bound molecules we imaged freely diffusing HaloTag. We found that only a small proportion of HaloTag molecules bind to chromatin (as observed previously^57,58^), and that many molecules have an anomalous exponent of around 1 [significantly higher than observed for fixed dye molecules (α of 0.62 ± 0.01) (**Extended Data Figure 6c**)]. We conclude that we can use 20 ms exposure trajectories to distinguish unconfined freely diffusing molecules from confined chromatin bound proteins. [We note, however, that the apparent diffusion coefficient of chromatin bound NuRD molecules (0.43 ± 0.03 μm^2^/s) cannot be distinguished from immobile when imaging using 20 ms exposures: we found that immobile dye molecules have a median precision of 60 nm and an apparent diffusion coefficient of 0.3 ± 0.2 μm^2^/s (**Extended Data Figure 6c**).]

### Chromatin association of the deacetylase sub-complex requires assembly into the intact holo-NuRD complex in ES cells

Having developed an approach to segment the 20 ms exposure trajectories of the NuRD complex into chromatin bound and freely diffusing molecules (**Figure 3a,b**), we investigated how the chromatin binding kinetics were affected by removal of MBD3, which disrupts the interaction between the HDAC- and CHD4-containing NuRD sub-complexes^18,24^ (**Figure 1a**).

**Figure 3.**
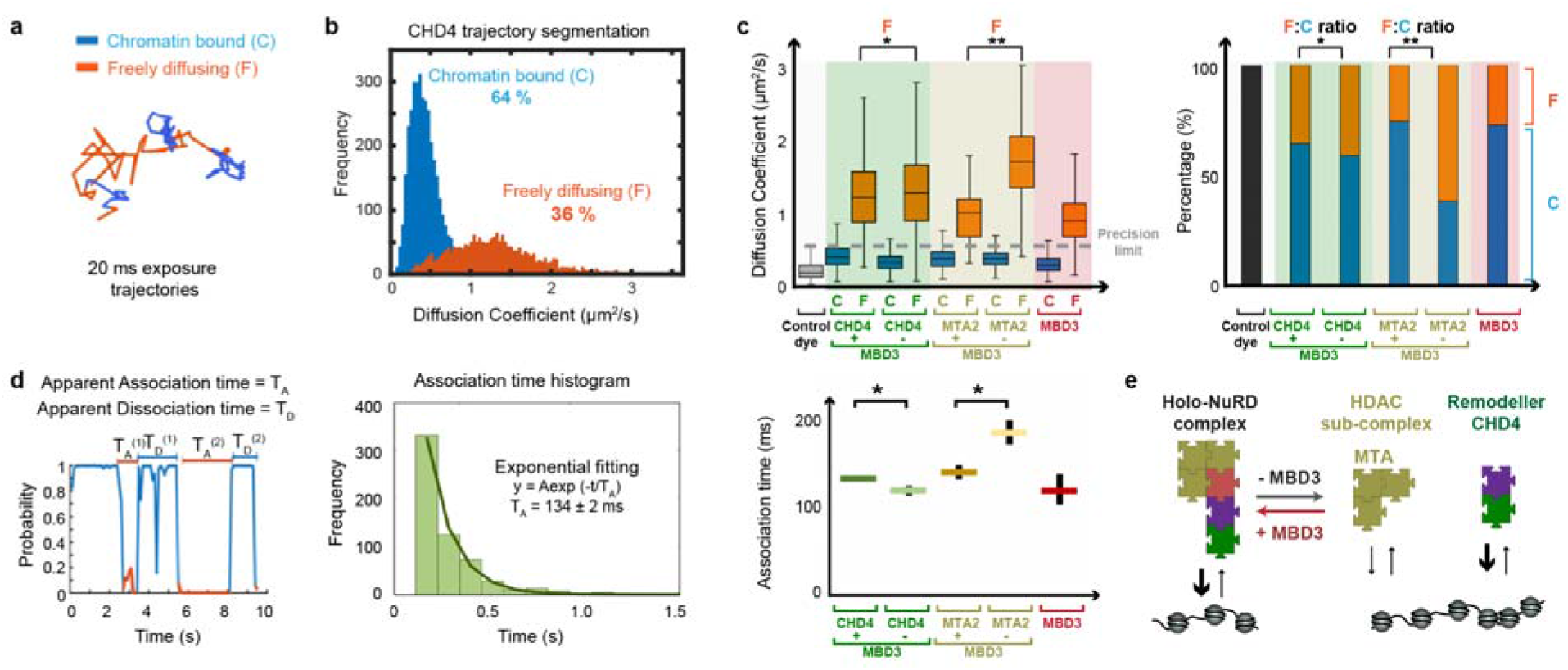
Live-cell single-molecule tracking reveals that the NuRD complex assembles before it binds to chromatin. (a) Segmentation of an example 20 ms trajectory into confined chromatin bound (C) (blue) and unconfined freely diffusing (F) states (orange). (b) Percentage of molecules and distribution of apparent diffusion coefficients for chromatin bound and freely diffusing CHD4 molecules. (c) (Left) Box plot of apparent diffusion coefficients for chromatin bound and freely diffusing CHD4 and MTA2 molecules in the presence and absence of MBD3 [*p < 0.01, **p < 0.001 (Kolmogorov-Smirnov test)]. The data for MBD3 is also shown as a control, and the grey dotted line indicates the upper bound of the precision limit calculated at the 95 % confidence interval for immobilised JF_549_ dye molecules. (Right) Percentage of freely diffusing and chromatin bound CHD4 and MTA2 molecules in the presence and absence of MBD3 [from Gaussian fitting, *p < 0.01, **p < 0.001 (2-way ANOVA)]. [The number of cells/trajectories used in the analysis were: 30/5,557 (CHD4), 25/2,337 (CHD4-MBD3), 10/336 (MTA2), 10/652 (MTA2-MBD3) and 30/2,224 (MBD3).] (d) (Left) A plot of the confinement probability allows determination of the association T_A_ and dissociation T_D_ times – defined respectively as the time a trajectory spends between periods of confined or unconfined motion. (Middle) A single exponential curve of rate lambda=1/T_A_ is then fit to the distribution of association/dissociation times. (Right) The association times extracted for CHD4 and MTA2 were then compared to those in the absence of MBD3, and to those for the MBD3 control [error bars show 95 % confidence intervals, *p < 0.01 (2-way ANOVA)]. (e) Schematic representation of a model in which MBD3-dependent assembly of the NuRD complex increases the association rate of the deacetylase sub-complex.

To explore whether the two sub-complexes are pre-assembled before binding to chromatin, we imaged the CHD4 remodeller and the HDAC-containing sub-complex (using tagged MTA2 molecules) in wild-type and *Mbd3*-ko ES cells. Single-molecule tracking of CHD4 in live ES cells revealed a small but significant increase in the apparent diffusion coefficient of freely diffusing CHD4 in the absence of MBD3 (1.05-fold, p = 0.009), consistent with the formation of smaller CHD4 sub-complexes^18,24,25^ (**Figure 3c**). [A larger increase might have been expected to arise from the disassembly of the holo-NuRD complex, but ES cells also contain CHD4 that is not present in NuRD (e.g. in the ChAHP complex with ADNP and HP1β,γ^59^). When we imaged the HDAC-containing sub-complex using tagged MTA2 molecules, this revealed a much more substantial increase in diffusion coefficient for freely diffusing MTA2 in the absence of MBD3 (1.7-fold, p < 10^−5^; **Figure 3c**), demonstrating that the deacetylase subcomplex is normally associated with CHD4 in the intact holo-NuRD complex (**Figure 1a**). [As a control, we also imaged tagged MBD3 and showed that both freely diffusing MBD3 and MTA2 molecules have similar diffusion coefficients, consistent with these two proteins associating with each other in the deacetylase sub-complex^18,24,25^ (**Figure 3c**).] Finally, we showed that MBD3 does indeed interact with CHD4 via GATAD2A, both *in vitro* using purified GATAD2A in pull-down reconstitution experiments (**Extended Data Figure 6b**), and in ES cells where knock-down of *Gatad2a* and *Gatad2b* increased the diffusion coefficient of CHD4 (1.05-fold, p < 10^−4^) (**Extended Data Figures 6d,e**).

We then examined how the NuRD complex interacts with chromatin by comparing the percentage of freely diffusing versus chromatin bound CHD4 and MTA2 molecules in the presence and absence of MBD3. We observed a small decrease in the percentage of CHD4 molecules bound to chromatin in the absence of MBD3, but there was a much more significant decrease (2.4-fold, p < 0.01) in the percentage of chromatin bound MTA2 molecules upon MBD3 depletion. This demonstrated that the chromatin bound deacetylase subcomplex is also normally associated with CHD4 in the intact holo-NuRD complex. It also suggested that CHD4 rather than the deacetylase sub-complex is primarily responsible for the association of NuRD with chromatin (**Figure 3c**). This finding was supported by *in vitro* experiments which showed that, in comparison to CHD4, the deacetylase subunit by itself does not bind strongly to nucleosomes (**Extended Data Figure 7a**). We conclude that NuRD normally exists as the intact complex *in vivo* and that, as with the removal of MBD3, depletion of GATAD2A/B is sufficient to unlink CHD4 from the HDAC-containing sub-complex in ES cells.

To investigate the chromatin binding kinetics of the CHD4 remodeller and the MTA2 deacetylase sub-complex in the presence and absence of MBD3, we next determined association times from the time spent freely diffusing between confined chromatin-bound states. We also attempted to determine dissociation times from the time spent bound to chromatin between unconfined freely diffusing states (**Figure 3d**). The distribution of association on times was well approximated by a single exponential, suggesting a Poissonian process. Consistent with our finding that CHD4 is primarily responsible for recruitment of NuRD to chromatin, we found no increase in the association time of CHD4 upon removal of MBD3. (Instead, we found a small decrease consistent with the faster diffusion of the smaller CHD4 sub-complex.) However, we did find a significant increase (1.3-fold, p < 0.01) in the association time of MTA2 upon MBD3 depletion (**Figure 3d**), consistent with the finding that the deacetylase sub-complex does not strongly associate with chromatin in the absence of CHD4 (**Figure 3c** and **Extended Data Figure 7a**). Although no changes in dissociation time were observed (**Extended Data Figure 7b**), we reasoned that our trajectories may be truncated by photobleaching. We therefore took advantage of ‘motion blurring’ when recording 500 ms trajectories to detect only chromatin bound proteins^1,60^, and combined this with time-lapse imaging using different interval lengths in an attempt to determine the dissociation time of the NuRD complex. To our surprise this showed that the dissociation times were much longer than we had expected (greater than 100 seconds for MBD3, **Extended Data Figure 7e**), such that it proved impossible to track individual molecules for long enough in order to determine reliable dissociation rates. We conclude that, once bound to a target site the intact NuRD complex binds for unexpectedly long times.

### The Holo-NuRD complex modulates chromatin movement at enhancers and promoters

Because chromatin bound NuRD cannot be distinguished from immobile dye when imaging using 20 ms exposures, we studied these slower moving molecules by imaging at a time resolution of 500 ms. (Here motion blurring also prevents imaging of freely diffusing molecules.) The 500 ms imaging was facilitated by the long residence times and trajectory lengths of the chromatin bound NuRD complex. Analysis of 500 ms exposure trajectories (that last more than 5 seconds) initially revealed two states of chromatin bound CHD4. Both the slow and fast states were primarily sub-diffusive (with an anomalous exponent α of around 0.5), and were characterised by different confinement lengths of 62 ± 12 nm and 110 ± 30 nm and apparent diffusion coefficients of 0.006 ± 0.002 and 0.018 ± 0.006 μm^2^/s (**Figure 2c**).

The apparent diffusion coefficient of the slow state was still below the precision limit even with 500 ms imaging: immobile dye molecules have a median precision of 34 nm and an apparent diffusion coefficient of 0.004 ± 0.003 μm^2^/s. Interestingly, however, visualisation of the fast state trajectories revealed a proportion of molecules exhibiting periods of directed motion, characterised by high anomalous exponents and high drift (**Figure 4a**). To determine whether there is indeed a proportion of molecules undergoing directed motion, we used Gaussian fitting to characterise the distributions in anomalous exponent for both the slow- and fast-states of chromatin bound CHD4 (in separate calculations, see **Extended Data Figures 8a-c**). This revealed a single slow state (S) with α_s_ of 0.59 ± 0.01 (67 % of sub-trajectories) and two fast states (F1 and F2) with different anomalous exponents: α_F1_ of 0.60 ± 0.01 (26 %) and α_F2_ of 0.89 ± 0.02 (7 %) (**Figure 4b**). Molecules in the fast F1 state have the same distribution of anomalous exponents as in the slow state and therefore explore the same chromatin environment. However, they diffuse faster and have a larger length of confinement and thus move further within the nucleus (**Figure 4b**). As can be seen in the example in **Figure 4a**, molecules in the fast F2 state, however, have a higher anomalous exponent and they explore a larger area of the nucleus than those in both the slow and the fast F1 states. Moreover, they have higher drift, indicative of movement in a directed manner: this is also consistent with the higher anomalous exponent.

**Figure 4.**
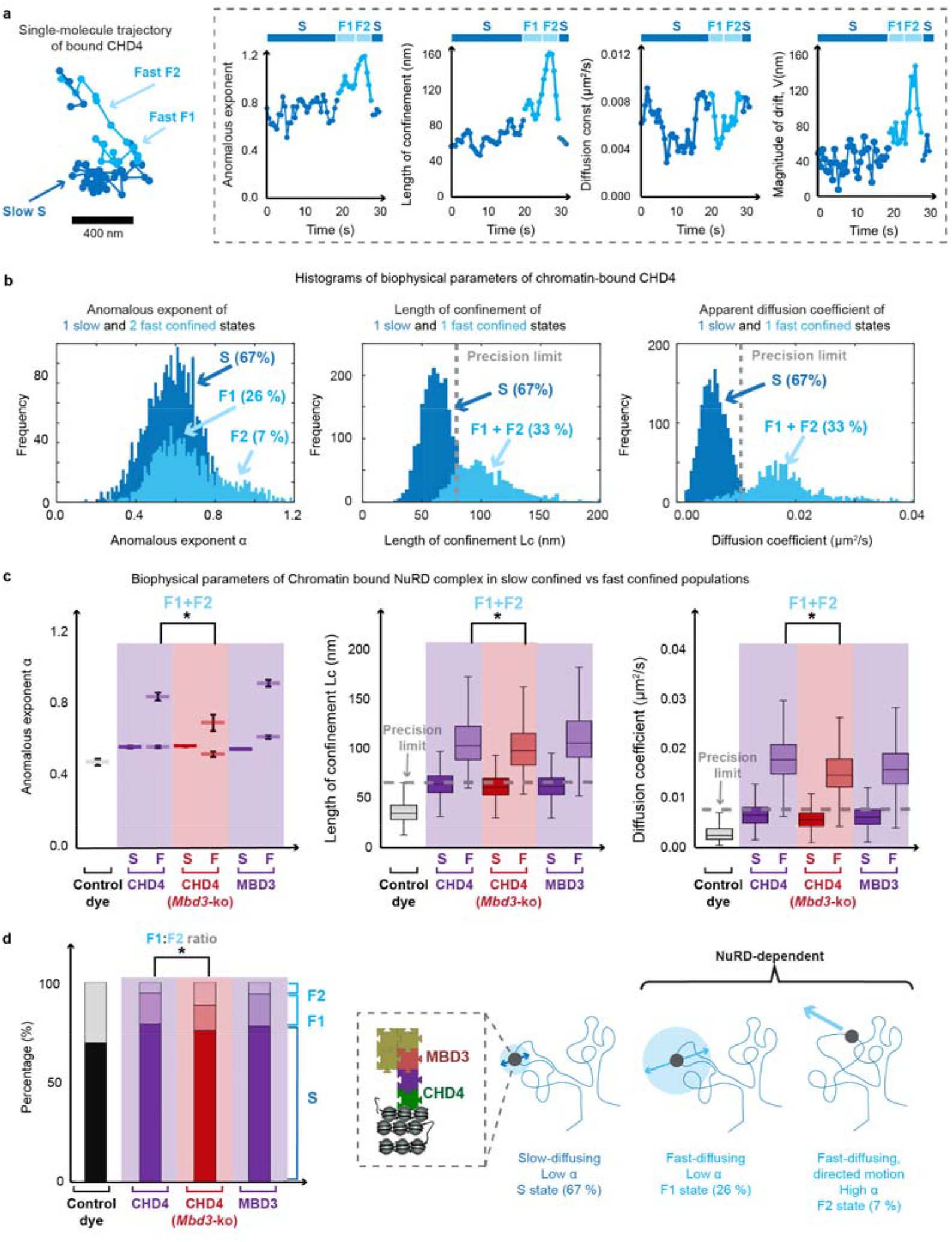
Assembly of the intact NuRD complex decondenses chromatin and reduces directed motion. (a) (Left) Example trajectory of a chromatin bound CHD4 molecule showing periods of both slow (dark blue) and fast (light blue) sub-diffusive motion. Two fast states (F1 and F2) are observed, with the F2 state showing periods of directed motion. (Right) The four biophysical parameters calculated along this trajectory with the fast F2 sub-trajectories showing a higher anomalous exponent, increased length of confinement and increased drift. (b) (Left) Gaussian fitting to the distribution of anomalous exponents identified a single α for slow confined NuRD complex molecules (dark blue), and two α values for molecules in the faster confined state (light blue). The resulting distribution of the lengths of confinement (Middle) and apparent diffusion coefficients (Right) are also shown. (c) Comparison of biophysical parameters for the CHD4 remodeller in the presence and absence of MBD3, and for MBD3 itself [the number of cells/trajectories used in the analysis were: 30/3,059 (CHD4), 15/2,111 (CHD4-MBD3), and 30/1,816 (MBD3)]. (Left) The anomalous exponents resulting from Gaussian fitting (error bars show 95 % confidence intervals, *p < 0.01, 2-way ANOVA); (Middle) Boxplots of the lengths of confinement; and (Right) the apparent diffusion coefficients are shown with the precision limit of the single molecule imaging indicated by a grey dotted line (*p < 0.01, Kolmogorov-Smirnov test). (We imaged JF_549_ dye fixed on a coverslip and found that all three biophysical parameters were significantly lower than the S, F1 and F2 states of NuRD components.) (d) (Left) Percentage of molecules in the slow or fast chromatin bound states (from the Gaussian fitting, *p < 0.01, 2-way ANOVA). (Right) Schematic representation of the three states of chromatin bound NuRD: a slow sub-diffusive S state with small exploration volume, one fast sub-diffusive F1 state with a larger length of confinement (exploration volume) and another fast sub-diffusive F2 state exhibiting directed motion.

We next compared the dynamics of chromatin bound MBD3 to that of CHD4 and found that it too exhibited the one slow and two fast states. Both chromatin bound MBD3 and CHD4 molecules exhibited motion in the fast F1 and F2 states in around 22-26% and 7-8 % of trajectories, respectively, confirming that these states are a property of the intact NuRD complex and not just of CHD4 (**Figures 4c,d**). Importantly, visualisation of the trajectories identified individual molecules that switch between the three states; S, F1 and F2 (**Figure 4a**). Thus, the three states are unlikely to either represent CHD4 forming different complexes or NuRD complex molecules bound in different regions of the nucleus.

We then compared the dynamics of chromatin bound CHD4 molecules in the *Mbd3*-ko and wild-type cells. Surprisingly, we found that the anomalous exponent, length of confinement and apparent diffusion coefficient of the fast F1 and F2 states were all higher in wild-type cells (**Figure 4c**). The increase in anomalous exponent in wild-type cells unexpectedly suggests that in the presence of the intact NuRD complex chromatin is less *condensed*, whilst the increased apparent diffusion coefficient and length of confinement show that CHD4 molecules diffuse more rapidly and explore a larger nuclear volume. We had expected to find that recruitment of the deacetylase by CHD4 would lead to less acetylated chromatin and *greater condensation*^61–65^ – and that the bound CHD4 molecules in wild-type cells would thus explore a smaller nuclear volume. In addition, in the presence of intact NuRD we also observed a significant decrease in the proportion of CHD4 molecules in the fast F2 state exhibiting directed motion (7.4 % in wild-type cells vs 18% in *Mbd3*-ko cells; see **Figure 4d** and **Extended Data Figure 8d**).

The fast F1 and F2 states of chromatin bound NuRD could result from movement on DNA due to chromatin remodelling or, bearing in mind the long dissociation times we determined for CHD4 (see above), from movement of NuRD-bound enhancers. To distinguish between these possibilities we targeted sites nearby active enhancers with dCas9-GFP, either by transfecting a published and validated CARGO vector expressing 36 different gRNAs targeting a *Tbx3* enhancer^3^ or by transfecting a single gRNA that targets DNA repeats near the *Nanog* gene (**Extended Data Figures 9a,b**). Targeting using our single gRNA was validated by confirming co-localisation of GFP-tagged dCas9 (detected by immunofluorescence) with DNA FISH probes (**Extended Data Figure 9c**). We carried out these experiments in cells that express an ER-MBD3-ER (estrogen receptor-MBD3-estrogen receptor) fusion protein in which the nuclear localisation of MBD3 (and thus assembly of the intact NuRD complex) is tamoxifen-inducible^18^. In this system the intact NuRD complex assembles and remodels chromatin/transcription factor binding following induction: after 24 hours the transcription factor landscape has been reset and transcriptional changes have occurred. This allowed us to study the new chromatin environment created by NuRD shortly after it had become established (thus allowing us to distinguish direct from downstream effects). We imaged cells showing bright undivided foci to avoid using data from cells in the S or G2 phases of the cell cycle, which exhibit blurred foci or doublets (**Extended Data Figure 10a)**. Because the background fluorescence from freely diffusing dCas9-GFP prevented 3D tracking of genomic loci on our DH-PSF microscope, we tracked the enhancer loci in 2D. Although this meant that we could not directly compare the parameters obtained in the 2D (active enhancer) and 3D (NuRD single molecule) tracking experiments, classification of the sub-trajectories once again revealed one slow and two fast chromatin states. The enhancer locus trajectories (**Figure 5a**) were also much longer than those of chromatin bound single CHD4 molecules (**Figure 4a**) allowing a better characterisation of switching between the different fast states (**Extended Data Figures 10b-d**). We found that, as when tracking CHD4 single molecules, addition of MBD3 (and thus intact NuRD assembly) significantly increased the apparent rate of diffusion of both enhancers in the fast-diffusing F1 and F2 states (**Figure 5b**). As when tracking CHD4, an increased proportion of sub-trajectories in the fast decondensed F2 state in wild-type cells was observed for both the *Tbx3* and *Nanog* enhancers (**Figure 5c**).

**Figure 5.**
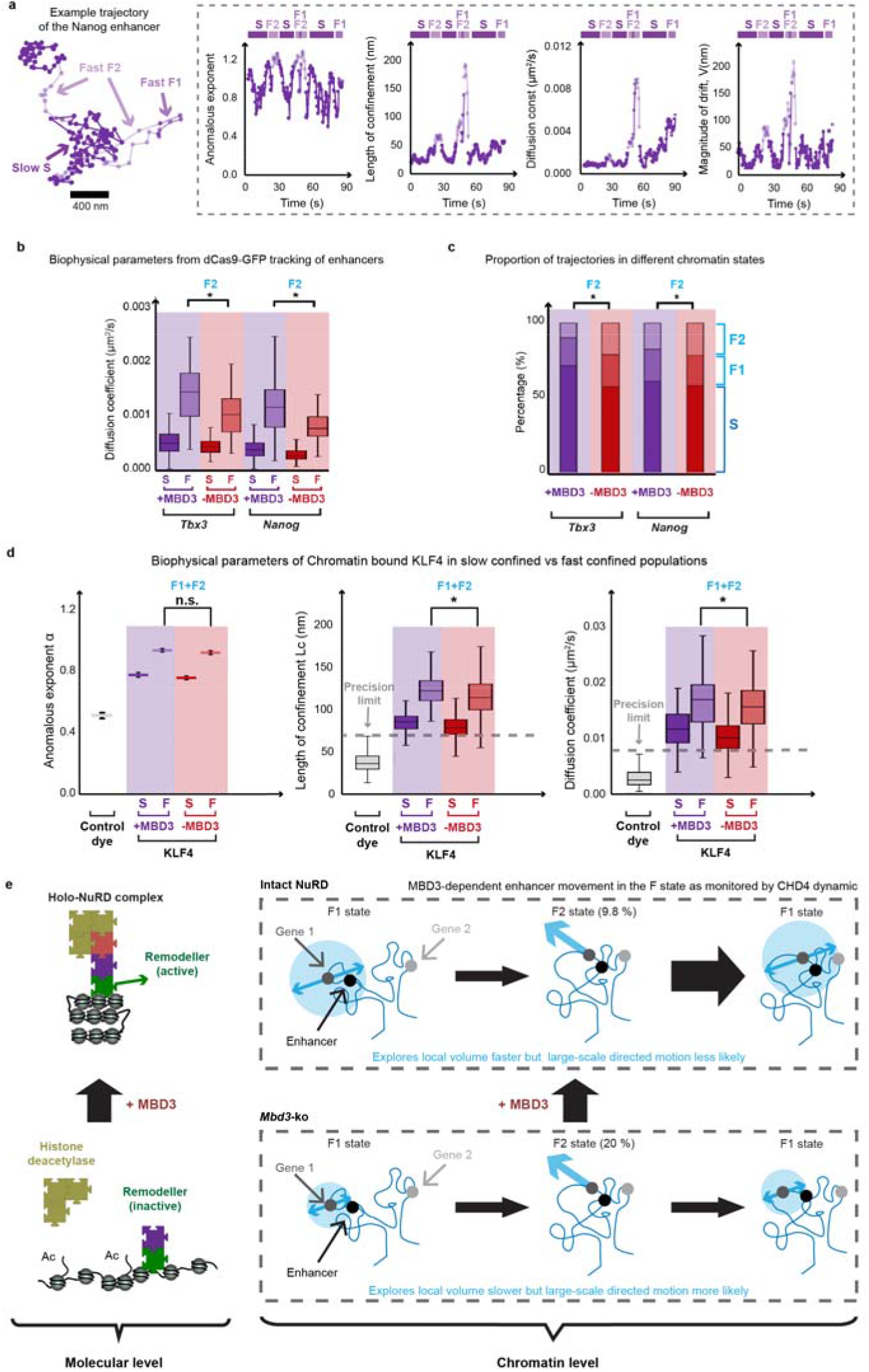
Assembly of the intact NuRD complex modulates the movement of active enhancers. (a) (Left) Example trajectory of the *Nanog* enhancer segmented to show periods of slow and fast sub-diffusive motion (dark and light blue respectively). (Right) The anomalous exponent, length of confinement, apparent diffusion coefficient and the magnitude of drift of the locus extracted from the trajectory shown. (b) Boxplots of the apparent diffusion coefficient calculated for 2D trajectories of loci nearby the *Tbx3* and *Nanog* enhancers, in the presence and absence of MBD3 (*p < 0.01, Kolmogorov-Smirnov test). Number of trajectories: 237 (*Tbx3*+MBD3) and 287 (*Tbx3*-MBD3); 546 (*Nanog*+MBD3) and 229 (*Nanog*-MBD3). (c) Gaussian fitting to the distribution of the anomalous exponents identifies a single slow and two faster states for enhancer loci. The percentage of sub-trajectories of enhancer loci exhibiting slow (S) and fast chromatin motions with either a low (F1) or high (F2) anomalous exponent α is shown (*p < 0.01, 2-way ANOVA). (d) Comparison of biophysical parameters for the KLF4 transcription factor in the presence and absence of MBD3. (Left) The anomalous exponents (error bars show 95 % confidence intervals from Gaussian fitting, *p < 0.01, 2-way ANOVA). (Middle and Right) Boxplots of the lengths of confinement and the apparent diffusion coefficients are shown with the precision limit of the single molecule imaging indicated by grey dotted lines (*p < 0.01, Kolmogorov-Smirnov test). Number of cells/trajectories used in the analysis were: 57/179 (KLF4+MBD3), and 56/652 (KLF4-MBD3). (e) Schematic model depicting: (Left) a molecular level view of how assembly of the NuRD complex affects chromatin remodelling^18^; and (Right) a chromatin level view of how intact NuRD increases the volume explored by an enhancer while at the same time reducing its likelihood of entering the F2 state in which the directed motion of chromatin might re-organise enhancer-promoter proximity.

To further confirm the effect of NuRD on enhancer dynamics we also imaged endogenous chromatin bound KLF4, a transcription factor which like NuRD binds to active enhancers in ES cells^18^ and which also clusters with NuRD in 3D genome structures^17^. We carried out 3D single molecule tracking of KLF4-HaloTag both before and 24 hours after inducing nuclear localisation of MBD3, and thus assembly of the intact NuRD complex, using tamoxifen. Once again we observed a significant increase in the length of confinement and apparent diffusion coefficient of both the slow and fast chromatin bound states of KLF4 (**Figure 5d**). The observation of similar effects 24 hrs after reassembly of the NuRD complex (and also at shorter times of 6 hrs after reassembly, data not shown), when either tracking active enhancers or chromatin bound KLF4, suggests that we are looking at a direct effect of NuRD on enhancer dynamics. Finally, to strengthen our conclusion that chromatin decompacts in the presence of the intact NuRD complex, leading to an increase in intermediate-range chromatin interactions, we carried out two-colour enhancer-promoter DNA FISH studies of three key pluripotency genes *Bmp4, Sox2* and *Tbx3*. In the presence of NuRD we found a significant increase (p<0.001) in the distance between their enhancers and promoters indicative of chromatin decompaction (**Extended Data Figures 9d,e**).

### Relationship of NuRD mediated modulation of enhancer dynamics to previous studies

How might the fast sub-diffusive states of the chromatin bound NuRD complex (F1 and F2) relate to the previous enhancer/promoter tracking experiments of Gu *et al*. (2018), and their transcription-dependent enhancer ‘stirring’ model? The similarity in both the diffusion parameters and anomalous exponents of their slow/fast and our S/F1 states, suggests that these correspond to either inactive (slow/S) or actively transcribed (fast/F1) enhancers and promoters. In our experiments, however, we additionally observe a fast F2 state which exhibits directed motion and we wondered whether this was due to chromatin movement resulting from transcriptional elongation? We therefore tracked bound CHD4 molecules after adding DRB, a small molecule inhibitor of transcriptional elongation^66^. Consistent with the results of Gu *et al*. (2018), premature termination by DRB led to a modest reduction in the proportion of bound CHD4 molecules exhibiting the fast F1 (transcription dependent) motion (from 26 % to 19 %). However, there was no reduction in the proportion of molecules in the fast F2 state nor change in the chromatin environment (i.e. in the anomalous exponent or length of confinement) in the presence of a block on transcriptional elongation (**Extended Data Figures 8b,d**). Finally, we tracked MBD3 molecules while blocking HDAC1/2 deacetylase activity with FK228^67^. As was the case when we prevented the association of the deacetylase sub-complex with CHD4 (see **Figure 4c**) we again observed a decrease in the anomalous exponent (reflecting a more condensed state), but there was no significant change in the proportion of molecules in the fast decondensed F2 state (**Extended Data Figures 8b,d**). We conclude that the fast F2 state is not dependent on transcriptional elongation or deacetylation activity. Instead, the results suggest that the assembly of the intact holo-NuRD complex (from both the chromatin remodelling and deacetylase subunits) modulates the dynamics of active enhancers, affecting their conversion between the slow S, fast F1 and fast F2 states (**Figure 5e**).

### The NuRD complex modulates the dynamics of enhancers to control transcription

Whilst we cannot rule out that some of the fast motions we observe when imaging the chromatin bound NuRD complex might result from chromatin remodelling, given that we observe similar dynamics for chromatin bound NuRD, active enhancers and chromatin bound KLF4, the results suggest that single molecule tracking of chromatin bound NuRD largely monitors the spatial dynamics of active enhancers. How might assembly of the deacetylase sub-complex with CHD4 to form intact NuRD influence transcription? Histone acetylation is thought to lead to a looser packing between nucleosomes and chromatin decompaction^61–65^, but here, by studying compaction in the immediate vicinity of the stably bound NuRD complex, we unexpectedly find that assembly of the intact NuRD complex leads to a more open chromatin structure where enhancers can diffuse faster and explore more of the nucleus (see **Figure 5e**). The chromatin decompaction and faster movement of enhancers in the presence of intact NuRD provides a molecular explanation for the increase in intermediate-range chromatin interactions (**Figure 1b**), the blurring of TAD and A/B compartment boundaries (**Figure 1c**), the increase in longer range enhancer-promoter interactions (**Figure 1d**), and the changes in enhancer interactions with promoters that are regulated by NuRD (**Figure 1e**). Our experiments also show that assembly of the intact NuRD complex reduces the time NuRD-bound enhancers spend undergoing directed motion, and we propose that this may limit the re-organisation of enhancer-promoter interactions leading to more stable, long-lived interactions. This could explain the observed increase in transcriptional noise, or low-level inappropriate transcription, observed in both human and mouse ES cells lacking functional NuRD^68,69^. CHD4 in the ChAHP complex has previously been shown to inhibit CTCF/Cohesin binding at repetitive DNA sequences^70^. Our results reveal that CHD4 in the NuRD complex plays a more general role by modulating CTCF/Cohesin interactions and cross-linking in active chromatin to promote an environment where enhancers and promoters can contact each other over longer distances, thereby reducing the fast F2 motions that our imaging approach has uncovered.

## Online Methods

### In-nucleus chromosome conformation capture (Hi-C)

#### Data acquisition

In-nucleus Hi-C was carried out as previously described^17^ on E14 wild-type and 7g9 *Mbd3-ko* (see Ref. 31 for construction and characterisation) ES cells. 50 bp paired-end sequencing was carried out on a HiSeq4000 instrument. Our own Hi-C analysis of *Mbd3-ko* ESCs cultured in 2i consisted of three experiments: SLX-18035, SLX-7676 and SLX-19611 (which was sequenced twice). Alongside these four datasets, we also collected wild-type 2i data (SLX-7672) and compared this with recently published Hi-C data for ESCs cultured in 2i conditions (https://www.ebi.ac.uk/arrayexpress/experiments/E-MTAB-6591/) (**Extended Data Figure 1a**). We used 4 replicates of this published data (ERR239137, ERR239139, ERR239141, ERR239143).

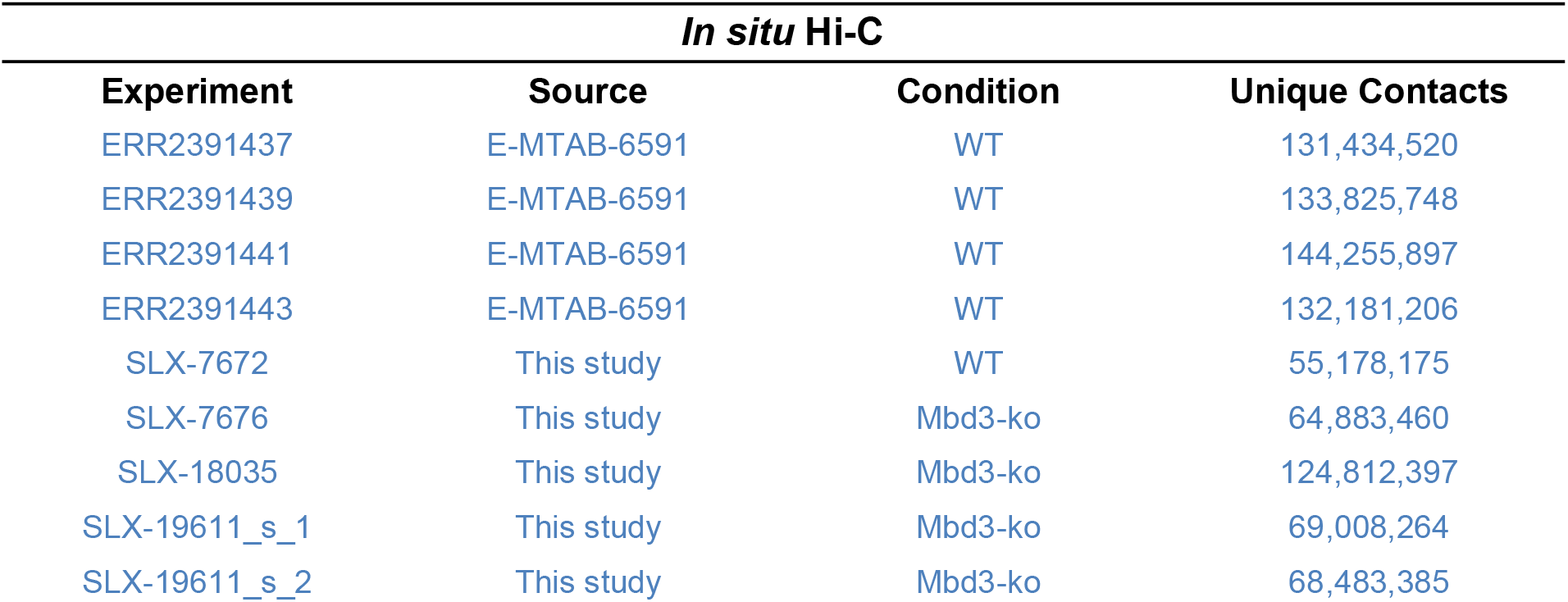

#### Pre-processing

For each experimental condition and replicate, FASTQ files were first processed and aligned against the mouse GRCm38.p6 reference genome using the nuc_processing package (https://github.com/tjs23/nuc_processing). The number of unique contacts observed varied significantly between replicates and conditions but overall a high number of reads were obtained in both conditions._The resulting raw contacts in NCC format were converted into hic and cooler format contact matrices using the juicer Pre (https://github.com/aidenlab/juicer/wiki/Pre) and HiCExplorer hicConvertFormat tools for downstream analysis (https://hicexplorer.readthedocs.io/en/latest/content/tools/hicConvertFormat.html)^71–73^. To assess the correlation between replicates within and between the two experimental conditions, we made use of the HiCExplorer hicPlotdistvscounts tool. Clustering of replicates based on contact distance distributions identified clusters for the two experimental conditions showing good agreement between replicates. We therefore merged the obtained contacts resulting in 705,063,441 and 327,180,884 contacts for the wild-type and *Mbd3-ko* conditions, respectively. Due to the difference in read coverage, we down-sampled our merged wild-type dataset to have the same number of contacts as our merged *Mbd3-ko* dataset. Finally, we performed Knight-Ruiz normalisation of the merged datasets to ensure balanced matrices.

#### Compartment Analysis

To identify A/B chromatin compartments within our merged datasets we made use of the recently developed CscoreTool (https://github.com/scoutzxb/CscoreTool)^74^. This tool assigns each genomic window *i* a score ∈ [−1,1] with =1 assigning a probability 1 that window *i* is in the A-compartment while = −1 assigns a probability 1 that window *i* is in the B-compartment. Importantly, unlike eigenvector analysis, c-scores of different samples can be directly compared since they represent probabilities. Binary compartments can also be assigned based on the sign of the c-score.

#### TAD Analysis

TAD calling was performed using Lavaburst, a recently published tool^41^ which uses an insulation-style metric called the TAD-separation score to identify the degree of separation between the up and downstream region at each Hi-C matrix bin. Local minima of the TAD separation score are considered as putative TAD boundaries and assigned q-values for calculation of a false-discovery-rate (fdr). For this analysis we used parameters gamma = 10 and beta = 50,000. For both datasets, we identified TADs using a binning resolution of 25 kb.

#### Enhancer-Promoter Links

We wanted to investigate whether transcriptional mis-regulation in *Mbd3-ko* ESCs was significantly associated with a disruption of cis-regulatory interactions. To identify putative promoter-regulatory element (RE) interactions we made use of a recent study^50^ which profiled RE-promoter interactions via thousands of separate CRISPR deletions. In particular, Fulco et al. (2019) found that a relatively simple activity-by-contact (ABC) model could be used to identify, with reasonable precision and recall, functional RE-promoter interactions. Crucially, the ABC model produced interactions with significantly higher accuracy than those identified using either linear distance along the genome or Hi-C contacts alone.

The ABC model considers the genome in 5 kb bins. For each promoter *p* and regulatory element *r* we define:

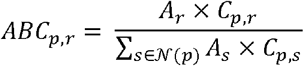

Where 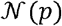 is the set of all regulatory elements within 5Mb of *p*, *C_p,r_* is the contact frequency between *p* and *r*, and *A_r_* is the activity of *r*. For regions of the genome with poor read coverage, *C_p,r_* is estimated assuming a power-law decay of contact frequencies:

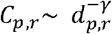

where *d_p,r_* is the genomic distance between the promoter and regulatory element while *γ* is inferred from the Hi-C contact maps. Fulco et al. (2019) define the activity of RE *r* as the geometric mean of read counts of DNaseI hypersensitive (DHS) and H3K27ac chromatin immunoprecipitation sequencing (ChIP-seq) at *r*. We did not have DHS tracks for our *Mbd3-ko* ES cells and therefore implemented our own version of the ABC model where we used the read counts in just the H3K27ac ChIP-seq data to score the activity of each regulatory element. Specifically, we identified putative regulatory elements using H3K4me3 and H3K27ac ChIP-seq data from wild-type (WT) cells and cells where MBD3 had been depleted^18^ (KO) as follows:

1. promoter regions were assigned as those regions ±1*kb* of a TSS and overlapping with an H3K4me3 ChIP-seq peak,
2. H3K27ac ChIP-seq peaks from that condition were considered as an initial putative list of condition specific RE’s,
3. H3K27ac peaks closer than 500bp were merged,
4. Peaks with total length <500bp were discarded,
5. Peaks overlapping with promoter regions were discarded, and
6. The union of the non-promoter peaks and promoter regions was then treated as a master list of putative RE’s which were each assigned a condition-specific score based on the mean H3K27ac peak strength across that RE.

In particular, REs could be assigned within a single condition as being either ‘intergenic’ or ‘promoter’ associated. Since REs were defined per condition, WT-unique, KO-unique and common overlapping peaks were assigned unique IDs for downstream analysis. The relevant code to perform this analysis and subsequent calculation of ABC scores can be found at: (https://github.com/dhall1995/Acitivity-By-Contact_Enhancer-Promoter_Link_Prediction).

Using calculated ABC scores, WT and KO links were assigned unique link-IDs based on the promoter and enhancer pair in question as well as the promoter genomic position in the case of a gene with multiple possible promoter regions. The top 10% of all identified links (WT & KO) when ranked by ABC score were then selected as ‘strong’ links. In our data this corresponded to an ABC threshold of ~0.12. Based on this threshold, unique link IDs could be assigned as either WT-unique, KO-unique or common depending on the conditions in which we observed that link. These thresholds were chosen to maximise the precision of identified links while retaining a significant number of links to analyse although we acknowledge that maximising precision comes at the expense of recall using the ABC model. Finally, after identification of WT-unique, KO-unique and common links, enrichment analysis was performed by associating each link with a promoter and performing χ^2^ analysis using the statsmodels python module.

### CTCF and SMC3 Cut&Run

Cut&Run experiments were carried out according to Meers *et al*. (2019)^75^ using a *Drosophila* genomic DNA spike-in and “input” controls consisting of samples which were processed in parallel with the Cut&Run samples but with untethered MNase. Western blots of nuclear lysates were also carried out as previously described^68^ to measure the relative levels of CTCF and SMC3 in wild-type and *Mbd3-ko* ES cells (**Extended Data Fig 4a**). The antibodies used were:

| ANTIBODY | RAISED IN | COMPANY | CATALOGUE NUMBER | WESTERN BLOT | ChIP | Clonality |
| --- | --- | --- | --- | --- | --- | --- |
| αCTCF | Rabbit | Millipore | 07-729 | 1:1000 | 5μl | Polyclonal |
| αSMC3 | Rabbit | Abcam | ab9263 | 1:1000 | 5μl (5μg) | Polyclonal |

50 bp paired-end sequencing was carried out on a Novaseq instrument – three biological replicates per sample were obtained with 8-16 million mapped reads/replicate, respectively, whilst the input samples had 8-23 million mapped reads/replicate, respectively. All Cut&Run data was trimmed using trim_galore and then aligned using standard Bowtie parameters to the Mus musculus reference genome (mm10) (https://www.bioinformatics.babraham.ac.uk/projects/trim_galore/). Heatmaps of Cut&Run enrichment were made using Deeptools v2.5.0^76^. Cut&Run bigwig tracks were calculated using BedTools and bedGraphToBigWig with standard parameters. The coverage was calculated with computeMatrix reference-point with options --binSize 10. The heatmap of standardised signal was then plotted using plotHeatmap. Peaks were called using MACS2 so as to give a False Discovery Rate of 1% and above 5-fold enrichment. All Venn Diagrams were plotted using the matplotlib-venn library in Python v3.6.

### Mouse embryonic stem cell line generation

Mouse embryonic stem (ES) cell lines were cultured in 2iL conditions^77^ – 50 % DMEM/F-12 medium (Gibco 21041025) and 50 % Neurobasal™ Medium (Gibco 12348017) supplemented with 1 x N2 to a final concentration of 2.5 μg/ml insulin (provided in-house by the Cambridge Stem Cell Institute), 0.5x B-27™ Supplement (Gibco 17504044), 1x minimum essential medium non-essential amino acids supplement (Sigma-Aldrich, M7145), 2mM L-glutamine (Life tech, 25030024), and 0.1 mM 2-mercaptoethanol (Life tech, cat. 21985023), 2i inhibitors (1 μM PD0325901, 3 μM CHIR99021) and 10 ng/ml mouse leukemia inhibitory factor (mLIF provided by the Biochemistry Department, University of Cambridge). Cells were passaged every two days by washing in PBS (Sigma-Aldrich, D8537), adding Trypsin-EDTA 0.25 % (Life tech, cat. 25200072) to detach the cells, and then washing in media before re-plating in fresh media. To help the cells attach to the surface, plates were incubated for 15 minutes at room temperature (RT) in PBS containing 0.1 % gelatin (Sigma Aldrich, G1890). All cell lines were routinely screened for mycoplasma contamination at least twice yearly and tested negative.

ES cells expressing CHD4 tagged at the C-terminus with HaloTag were generated in the presence and absence of MBD3, as previously described^17,24,25^. Briefly, this was achieved by CRISPR/Cas9 based knock-in of a cassette containing mEos3.2-HaloTag-FLAG and a puromycin selection gene into one *CHD4* allele. The puromycin cassette was then removed using Dre recombinase to generate the *CHD4* allele with a C-terminal HaloTag fusion. Since knockout of *CHD4* is lethal, we used cell viability assays and the ability to immunoprecipitate NuRD component proteins (**Extended Data Figure 4c**) to verify that the function of the tagged CHD4 was not significantly impaired. We similarly generated knock-in ES cells in an E14Tg2a (XY) background expressing MBD3 tagged at the C-terminus with HaloTag (**Extended Data Figure 4b**). MTA2-HaloTag knock-in cell lines were generated in MBD3-inducible ES cells^18^ (**Extended Data Figure 4d**), in which MBD3 is fused to the oestrogen receptor at both the N- and C-termini so that it initially localises at the cytoplasm but then translocates to the nucleus when induced with 4-hydroxytamoxifen added directly to the culture media to a final concentration of 0.4 nM. Western blots were carried out using nuclear lysates as previously described^18^, to confirm the expression and assembly of the NuRD complex (**Extended Data Figure 4a**). Immunoprecipitations were carried out using antibodies to CHD4 or MTA2, or Halo-Trap beads (ChromoTek) in the case of the CHD4-Halo line. The antibodies used were:

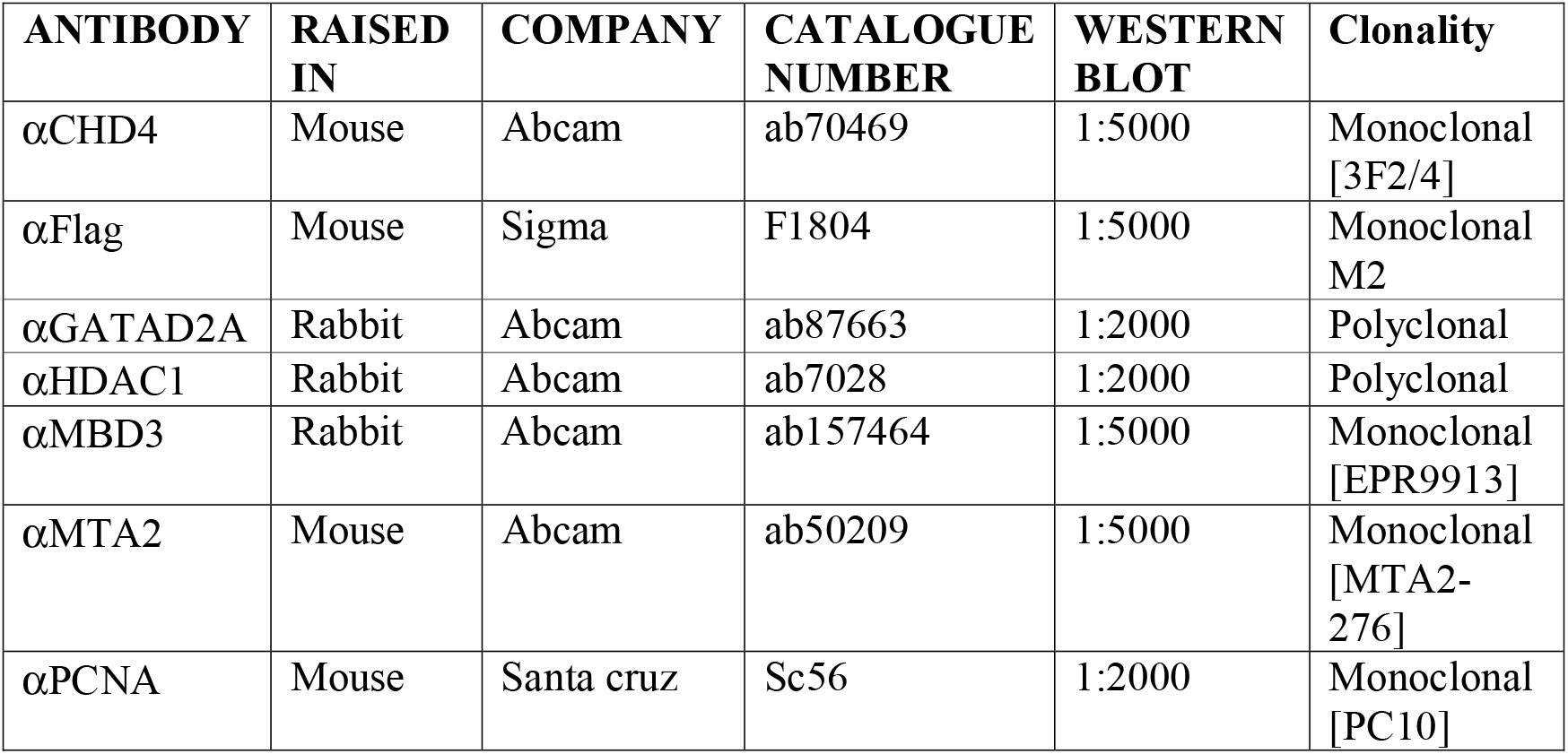

### Live-cell 3D single-molecule imaging

ES cells expressing HaloTag-tagged CHD4, MBD3 and MTA2 were passaged two days before imaging onto 35 mm glass bottom dishes No 1.0 (MatTek Corporation P35G-1.0-14-C Case) in serum/LIF imaging medium: Fluorobrite™ DMEM (Thermo Fisher Scientific, A1896701) containing 100 mM 2-mercaptoethanol (Life tech, cat. 21985023), 1x minimum essential medium non-essential amino acids (Sigma-Aldrich, M7145), 2 mM L-glutamine (Life tech, cat. 25030024), 1 mM sodium pyruvate (Sigma-Aldrich, S8636-100ML), 10 % fetal bovine serum (HyClone FBS, Lot nr SZB20006, GE Healthcare Austria SV30180.03) and 10 ng/ml mLIF (provided by the Biochemistry Department, University of Cambridge). Glass bottom dishes had been prepared by incubation in 0.01% poly-L-ornithine (Sigma-Aldrich P4957) for 30 minutes, followed by two rinses in PBS and incubation in PBS containing 10 μg/mL laminin (Sigma-Aldrich L2020) for at least 4 hours. Just before single-molecule imaging experiments, cells were labelled with 0.5 nM HaloTag^®^-JF_549_ ligand for 15 minutes, followed by two washes in PBS and a 30 minute incubation at 37 °C in imaging medium, before imaging the cells in fresh serum/LIF imaging medium. Cells were under-labelled to prevent overlap of fluorophores during single-molecule tracking experiments. The HaloTag dyes were a kind gift from Luke D. Lavis (HHMI).

For the HaloTag-NLS control, the pEF-HaloTag-NLS vector was generated by replacing the HP1 sequence in a previously described HaloTag-HP1 expression vector^78^ with a SV40 NLS sequence. 1μg of the pEF-HaloTag-NLS vector was transfected into ES cells using Lipofectamine 2000 Transfection Reagent (Thermo Fisher Scientific 11668019) during passaging onto 35 mm glass bottom dishes No 1.0 (MatTek Corporation P35G-1.0-14-C Case). Media was changed after 24 hours and samples were both labelled as above and imaged the following day.

Transcription elongation was inhibited using 100 μM 5,6-dichloro-1-β-D-ribofuranosylbenzimidazole (DRB) and deacetylase activity using 10 nM FK228 (TOCRIS Bioscience, UK) both for two hours prior to imaging^67,79^. qPCR experiments confirmed inhibition of transcription upon addition of DRB (data not shown).

A custom-made double-helix point spread function (DHPSF) microscope was then used for 3D single-molecule tracking as previously described^25^. The setup incorporates an index-matched 1.2 NA water immersion objective lens (Plan Apo VC 60×, Nikon, Tokyo, Japan) to facilitate imaging above the coverslip surface. The DHPSF transformation was achieved by the use of a 580 nm optimized double-helix phase mask (PM) (DoubleHelix, Boulder, CO, USA) placed in the Fourier domain of the emission path of a fluorescence microscope (Eclipse Ti-U, Nikon). The objective lens was mounted onto a scanning piezo stage (P-726 PIFOC, PI, Karlsruhe, Germany) to calibrate the rotation rate of the DHPSF. A 4*f* system of lenses placed at the image plane relayed the image onto an EMCCD detector (Evolve Delta 512, Photometrics, Tucson, AZ, USA). Excitation and activation illumination was provided by 561 nm (200 mW, Cobolt Jive 100, Cobolt, Solna, Sweden) and 405 nm (120 mW, iBeam smart-405-s, Toptica, Munich, Germany) lasers, respectively, that were circularly polarized, collimated, and focused to the back focal plane of the objective lens. Oblique-angle illumination imaging was achieved by aligning the laser off axis such that the emergent beam at the sample interface was near-collimated and incident at an angle less than the critical angle (θ_c_ ~ 67°) for a glass/water interface. The fluorescence signal was then separated from the excitation beams into the emission path by a quad-band dichroic mirror (Di01-R405/488/561/635-25x36, Semrock, Rochester, NY) before being focused into the image plane by a tube lens. Finally, long-pass and band-pass filters (BLP02-561R-25 and FF01-580/14-25, respectively; Semrock) placed immediately before the camera isolated the fluorescence emission. Using 561 nm excitation, fluorescence images were collected as movies of 60,000 frames at 20 ms or 4,000 frames at 500 ms exposure. A continuous 561 nm excitation beam at ~1 kW/cm^2^ was used for 20 ms exposure imaging and at ~40 W/cm^2^ for 500 ms exposure imaging. Each experiment was carried out with at least 3 biological replicates (3 fields of view, each containing around 3 cells).

### Residence time analysis from time-lapse 500 ms exposure imaging

Since photobleaching is related to the number of exposures, and the residence time is related to the time a molecule spends bound to chromatin, it is possible to change the time-lapse between exposures and use the data to extract both the residence time^1^ and photobleaching rate. However, when we imaged at time intervals of 0.5 s, 2.5 s, 8 s and 32 s, we discovered that at the longest time lapse (32 s) we could see no decrease in the mean number of frames imaged before photobleaching, implying that the residence time had no impact on the measurement, which was thus dominated by photobleaching (**Extended Data Figure 7e**). To estimate the residence time would likely require imaging at much longer time-lapses, but because chromosomes and the cell itself move significantly during periods longer than this, it becomes unreliable to track individual chromatin bound NuRD complex subunits.

### 3D single-molecule image processing, generation of trajectories and determination of experimental precision

Single molecules were localized from 3D movies using the easy-DHPSF software^80^ with a relative localization threshold of 100 for all 6 angles for the 20 ms data and relative thresholds of 116, 127, 119, 99, 73 and 92 for the 500 ms data. The trajectories of individual molecules were then assembled using custom Python code for connecting localisations in subsequent frames if they were within 800 nm for 20 ms trajectories and within 500 nm for 500 ms trajectories (https://github.com/wb104/trajectory-analysis). This code also outputs average signal intensity per trajectory and trajectory lengths (OPTION - savePositionsFramesIntensities) and a summary of this data is reported in **Extended Data Figure 7d**.

Our precision values (measured for fixed dye molecules on the coverslip and calculated using the approach described by Endesfelder *et al*. ^81^ were 60 nm and 34 nm for the 20 and 500 ms tracking experiments, respectively. The lower limits of the effective diffusion coefficient one can measure are dependent on the precision values. The effective diffusion coefficient (D_eff_) is equal to the displacement squared/time. Thus, if the upper limit of the precision is 60 nm, then the lower limit of D_eff_ that we can measure for 20 ms imaging is: 0.06*0.06/0.02 = 0.18 μm^2^s^−1^. For 500 ms imaging on the other hand, the upper limit of our precision was 34 nm, corresponding to a lower limit of D_eff_ that we can measure of: 0.034*0.034/0.5 = 0.002 μm^2^s^−1^. Consistently, when we measured the apparent diffusion coefficients for dye molecules attached to a coverslip we determined values of 0.3 ± 0.2 μm^2^/s and 0.004 ± 0.003 μm^2^/s, respectively (see **Extended Data Figures 6c and 8b**). Any measured diffusion coefficients below these values do not have a biophysical interpretation.

### Single-molecule trajectory analysis

We describe below (see Section 1) a switching model for a stochastic process between two or more states, characterized by different diffusion coefficients. We initially used this approach to decide whether we should develop a two- or three-state diffusion model to classify the distribution of displacements measured along single molecule trajectories. We then went on to develop an improved and novel algorithm to classify sub-trajectories into confined and unconfined states based on four physical parameters using a Gaussian mixture model. This is described in Section 2. The algorithm was then tested on simulated trajectories which provided ground truth data. Finally, in Section 3, we describe how we applied this classification algorithm to single particle trajectories to estimate the diffusion coefficients in confined and unconfined states. However, the original classification based on diffusion coefficients provided a complementary analysis, and justified the results we obtained later, which is why it is included here.

#### 1. Switching dynamics and displacement analysis

We describe the motion of a particle that switches between different states characterized by different diffusion coefficients^82,83^. We initially used this description to identify the most appropriate number of switching states, based on the distribution of the length of the instantaneous displacements Δ*X* = *X*(*t*+Δ*t*)−*X*(*t*). First, we describe the switching dynamics and fitting procedure, and then how we used the Bayesian Information Criterion (BIC) to select the optimal model.

##### 1.1 Stochastic motion described by a diffusion model

The dynamics of a chromatin-binding protein complex such as NuRD, or some component of it, can be described by Langevin’s equation^84,85^ in the classical overdamped limit, where the position *X*(*t*) satisfies:

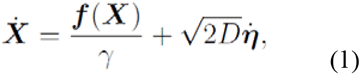

where the force *f* depends on the position *X* and the second term corresponds to steady-state diffusion in a crowded medium, characterized by a constant diffusion coefficient *D*. Here *η* is a Gaussian variable with mean 0 and variance 1. When the protein binds to its molecular partners or chromatin, the diffusion coefficient could change, and the motion can become restricted, e.g. to that along the path of the DNA.

Experimental trajectories consist of a series of points (*X*(*k*Δ*t*) *k* = 1… and in the absence of any additional localization error noise, or an external force *f* = 0, the displacement dynamics at the sampling rate interval Δ*t*, is given by:

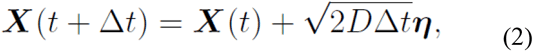

where *η* is a vector of Gaussian values of mean 0 and variance 1. In two dimensions, the distribution of displacements is:

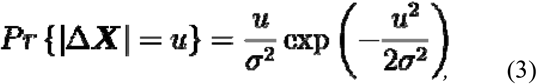

where *σ*^2^ = 2*D*Δ*t* (see Eq. 5). For empirical displacements that contain a Gaussian localization erroror,, characterized at the time resolution Δ*t* by the amplitude σ_le_, the formula should be modified^86^:

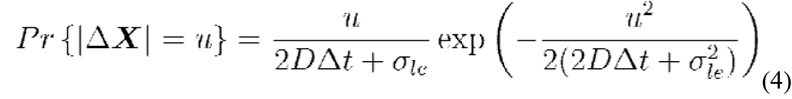

We can thus use the approximated diffusion coefficient:

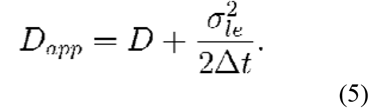

##### 1.2 Modelling stochastic switching behaviour

We next constructed a model for a chromatin-binding protein by using a classical Markov chain with two states *1* and *2*, characterized by rate constants *λ* and μ for switching between the states, and by the diffusion coefficients *D*_1_ and *D*_2._ The associated jump process is defined by:

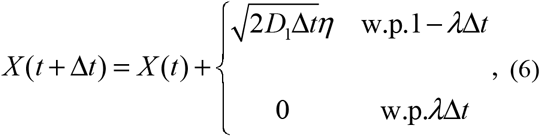

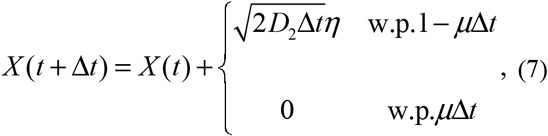

where the transition rates from states 1 to 2 are described by the switching equation:

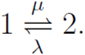

The probability density function (pdf) *p_d_* for the displacement of a molecule depends on the state of the process and can be computed using Bayes’ law:

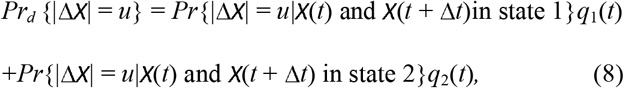

where using Bayes relation, for state *i* = 1 or 2,

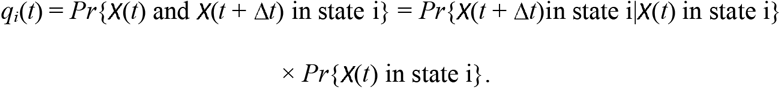

Because *p_i_*(*t*) = *Pr*{*X*(*t*) in state i} is the solution to the Master equation:

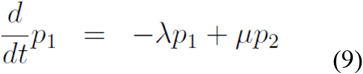

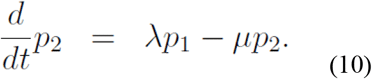

and

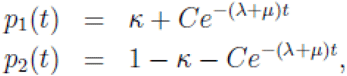

where 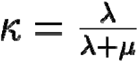 and *C* depend on the initial distribution, we have:

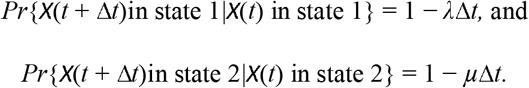

We conclude that in two-dimensions, for a long-time *t* and a short-time Δ*t*, using Eqs. 8 and 3, the pdf can be be written as:

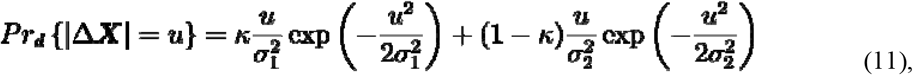

where *σ_k_* = 2*D_k_*Δ*t* and *k* = 1,2, and *κ* is the ratio of the time spent in a confined state. In the case of a three-state model, the pdf for the displacement of a molecule is:

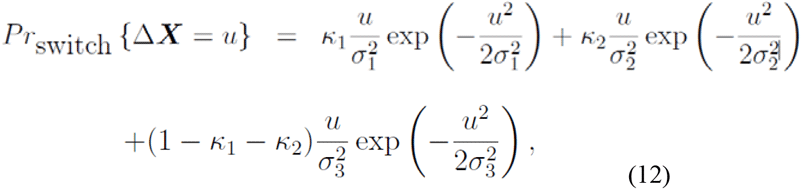

where *σ_k_* = 2*D_k_*Δ*t* with *k* = 1, 2, and 3 and *D*_1_, *D*_2_, and *D*_3_ are the three diffusion coefficients. Here *κ*_1_ (respectively *κ*_2_) is the fraction of time spent in state 1 (respectively state 2).

To test the appropriateness of a one-, two- or three-state models, we extracted the diffusion coefficients from Eqs. 11–12 and the associated rate constants. We then estimated the quality of fit of the different models by using the Bayesian Information Criterion (BIC) defined as:

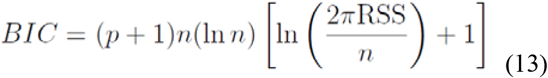

where *p* is the number of parameters of the model from Eq. 12, *n* is the number of data points, and *RSS* is the residual sum of squares between the model and the data.

Analysis of the 20 ms trajectories revealed that the data could be characterized by either a two or a three state model, consisting of a confined and either one or two unconfined states (see Table 1, below). Similarly, we evaluated 1, 2 and 3 state models for analysis of the 500 ms data consisting of a confined and either one or two sub-diffusive states (see Table 2, below). Although this showed that a three-state model could best describe both the 20 and 500 ms data, the analysis also revealed that the BIC values for the three-state model were only slightly larger, suggesting that a two-state model would be sufficient to explain the data.

**Table 1.**
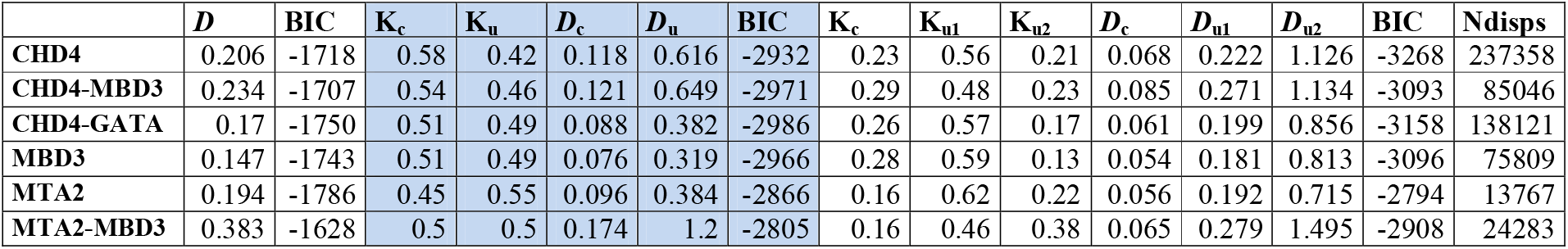
Parameter estimation from trajectories with Δ*t* = 20 ms.

**Table 2.** Parameter estimation from trajectories with Δ*t* = 500 ms. In both Tables 1 and 2, *D*_c_ and *D*_u_ are the estimated apparent diffusion coefficients (whilst K_c_ and K_u_ are their relative proportions). The blue boxes indicate our preferred model based on BIC analysis.

| | $D$ | BIC | $K_c$ | $K_u$ | $D_c$ | $D_u$ | BIC | $K_c$ | $K_{u1}$ | $K_{u2}$ | $D_c$ | $D_{u1}$ | $D_{u2}$ | BIC | Ndisps |
| --- | --- | --- | --- | --- | --- | --- | --- | --- | --- | --- | --- | --- | --- | --- | --- |
| CHD4 | 0.005 | -1874 | 0.55 | 0.45 | 0.003 | 0.01 | -3089 | 0.16 | 0.68 | 0.16 | 0.002 | 0.005 | 0.017 | -3170 | 91470 |
| CHD4+DRB | 0.003 | -1796 | 0.46 | 0.54 | 0.002 | 0.006 | -3036 | 0.24 | 0.63 | 0.13 | 0.001 | 0.004 | 0.014 | -3062 | 34164 |
| CHD4-MBD3 | 0.004 | -1855 | 0.52 | 0.48 | 0.002 | 0.008 | -3145 | 0.2 | 0.63 | 0.17 | 0.002 | 0.004 | 0.014 | -3218 | 60964 |
| MBD3 | 0.004 | -1827 | 0.55 | 0.45 | 0.003 | 0.009 | -3094 | 0.22 | 0.62 | 0.16 | 0.002 | 0.005 | 0.017 | -3205 | 53176 |
| MBD3+FK228 | 0.004 | -1803 | 0.56 | 0.44 | 0.002 | 0.008 | -3151 | 0.36 | 0.51 | 0.13 | 0.002 | 0.005 | 0.017 | -3181 | 64328 |

In summary, the displacement analysis revealed that a two-state diffusion model should be appropriate to analyse NuRD complex dynamics: in both cases there was only a small improvement in the BIC for the three-state diffusion model, suggesting that the 20 ms trajectories can be characterised by confined (low diffusion coefficient D) and unconfined (high diffusion coefficient D) states, whilst the 500 ms data can be characterised by slow and fast sub-diffusive chromatin bound states. However, this analysis supported our later identification of a slow and two fast states in the 500 ms data (see Section 5, below).

#### 2. Gaussian mixture model for classification and segmentation of trajectories into two states using four biophysical parameters (4P)

The Gaussian classification described in Section 1 above, is limited as it considers each displacement as being independent and then pools them, destroying any possible causality present in the trajectories. Moreover, because parts of our trajectories could correspond to molecules that are confined (i.e. chromatin bound) and other parts to when they are unconfined (or diffusing more freely), our trajectories exhibit a variety of behaviours characterised not only by slow and fast diffusion coefficients but also by smaller/larger anomalous exponents, the exploration of smaller/larger regions of space, and less/more directional movement. We therefore needed an algorithm that took into account these types of behaviour and not just changes in diffusion coefficient. Based on recent work to characterise such biophysical parameters^55^, we present here a new algorithm that uses these parameters to classify and segment a trajectory into two states – when analysing 20 ms data the resulting sub-trajectories are confined (C) and unconfined (U) in the associated four dimensional space, whilst for the 500 ms data, we observe two confined states, slow- and fast-diffusing (see **Figure 2** in the main text). The method is based on a generalized Gaussian mixture model^87^, where the input data is an ensemble of trajectories and the output is two ensembles of sub-trajectories.

##### 2.1 Input data

We obtain an ensemble of *N* trajectories *X_i_*(*k*Δ*t*), *i* = 1,…*N, k* = 0,1,…,*n_i_*, with an acquisition time step Δ*t*, such that each trajectory consists of *n_i_* discrete points in three dimensions.

##### 2.2 Statistical features are extracted in a sliding window along the trajectories

To classify sub-trajectories as confined C or unconfined U, we used four physical parameters^53,88^, computed along single trajectories. For a trajectory given by the successive points *X_i_*(*k*Δ*t*), we used a sliding window *W_k_* containing 2*l* + 1 points, centered at *X*(*k*Δ*t*), and defined as:

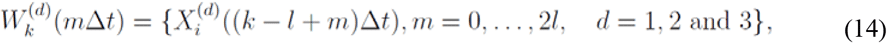

where *d* is the space dimension. The sliding window *W_k_*(*m*Δ*t*) is applied for *k* = 1,…*n_i_* along a trajectory containing *n_i_* points.

We now briefly discuss the four parameters we used for the classification based on previous work which showed that they can be used individually for the classification of single-molecule trajectories:

###### Anomalous exponent *α*

The anomalous exponent characterizes the motion of a stochastic particle on a particular time scale involving several time steps Δ*t*. It is computed from the Mean-Square-Displacement (MSD) 〈|*X*(*t* + Δ*t*) − *X*(*t*)|^2^〉 that behaves like Δ*t^α^* where Δ*t* is small compared to the time of the process. A value of *α* = 1 reflects Brownian motion, and an *α* > 1 is called super-diffusion, which may represent dynamics containing an element of deterministic (ballistic) directed motion. Finally, an *α* < 1 is sub-diffusive motion and has been used previously to study changes in chromatin compaction^53,89,90^. To estimate *α* for each point *X*, we first compute the MSD *S_k_* over the sliding window *W_k_*(*m*Δ*t*) classically defined by:

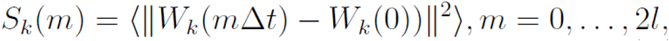

Where 〈·〉 denotes the average where we use the intermediate point from 1 to m. To estimate the exponent *α*_*i*_(*k*) for point *X*(*k*Δ*t*), we fitted the function *S_k_*(*m*) computed by summing over each displacement contributing to Eq. (15) with:

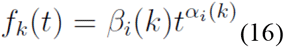

where *β*_*i*_(*k*) > 0. For the fit, we constrain the variable *t* in the ensemble [0,Δ*t*,…,(2*l*+1)Δ*t*].

###### Effective Diffusion coefficient D

We estimate the effective diffusion coefficient^88,91^ by computing the second statistical moment along the trajectories. We use the empirical estimator to estimate for each sliding window *W_k_*(*m*Δ*t*)^53^:

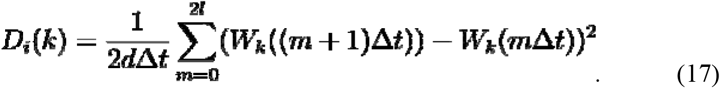

###### Length of confinement *Lc*

The length of confinement estimates the size of a domain where a trajectory is confined. It is computed empirically in the window *W_k_* by:

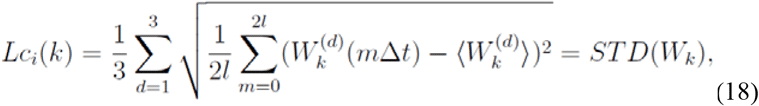

where:

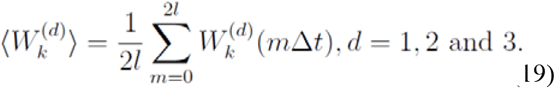

It is the standard deviation of the sub-trajectory *W_k_*, where the average position is 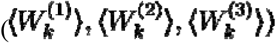.

###### Magnitude of the drift vector *V_i_* (k)

To characterize the displacement of a trajectory between the beginning and the end of the sliding window *W_k_*, we compute the magnitude *V_i_* (k) of the drift vector *V_i_* for each dimension *d* = 1,2,3, ^91^ using the formula:

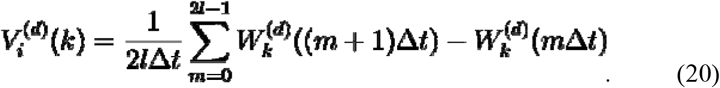

The norm of the drift *V_i_* (k) is:

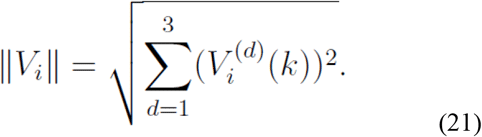

In summary, we compute four parameters: the anomalous exponent *α*, the apparent diffusion coefficient *D*, the length of confinement *Lc*, and the magnitude of the drift vector *V_i_* (k). These are all computed for each point along a trajectory using a sliding window. The sliding window of 11 points was chosen by trial and error. Below this value, the anomalous exponent distributions for confined and unconfined molecules tended to merge suggesting that long trajectories are essential for reliable estimation of this parameter. Above this value, the diffusion coefficient histograms for confined and unconfined molecules tended to merge suggesting that transitions between these populations occur leading to averaging of the diffusion coefficients. In the next section, we use these four parameters for classification.

##### 2.3 Classification of a discrete time point in a trajectory into either a confined (C) or an unconfined (U) state

To classify each time point *X*((*k* − 1)Δ*t*) for *k* = 1,…,*n_i_* of all trajectories indexed by *i* = 1,…,*N* into C and U classes, we first collected all the values from Eqs. 16–21, computed from each sliding window *W_k_* (Eq. 14). This led to a total of *n_i_* × 4 parameters, that we organized in a matrix *R_i_* associated with trajectory *i*:

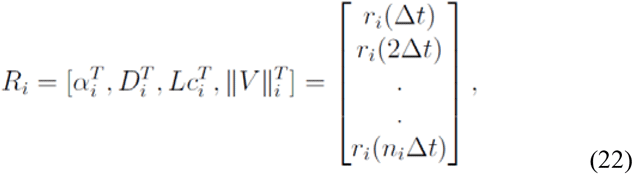

where *T* is the transpose operator, and *r_i_*(*k*Δ*t*) are four dimensional vectors of the parameters (Eqs. 16–18, 21).

We then concatenate all feature matrices *R_i_* (Eq. 22), *i* = 1,…*N* into the 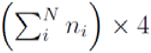 general matrices ***R***, defined by:

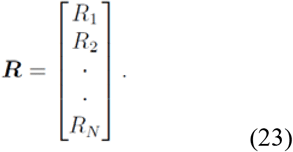

and we normalize each column (feature) in R^(*j*)^,*j* = 1,…,4 by subtracting its mean and dividing by its standard-deviation.

To separate the histograms of the four parameters into two independent classes, we constructed an unsupervised binary classifier using a two-component Gaussian mixture model in a four-dimensional space, corresponding to the four parameters Eqs. 16–21. The Gaussian mixture distribution *p* is a weighted sum of two multivariate Gaussian densities, defined as:

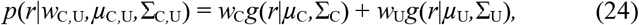

where *w*_C_,*w*_U_ are the mixing weights, such that *w*_C_ + *w*_U_ = 1, and *g*(*r*|*μ*,Σ) are the Gaussian densities

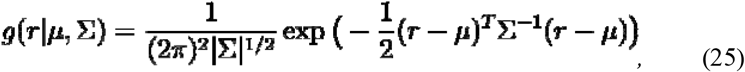

and Σ_C_,Σ_U_ and *μ*_C_,*μ*_U_ are the four-dimensional covariance matrices and mean vectors for components C and U, respectively. We then find the values of the parameters *μ*_C_,*μ*_U_,Σ_C_,Σ_U_ and *w*_C_,*w*_U_ of Eq. 24 which best separate the data as the maximal likelihood estimators of the density (Eq. 25), given the observed statistics in *R* (Eq. 23), by using the Expectation-Maximization algorithm^92^.

For each point *X_i_*(*k*Δ*t*), with its associated feature vector *r_i_*(*k*Δ*t*) (Eq. 23), we assign a label *n* ∈ {C,U}, based on the posterior probability *P* of the density Eq. 25, given by:

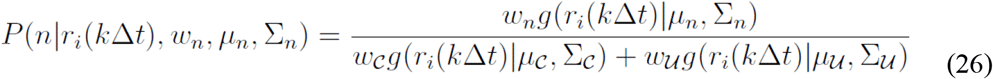

such that for each point:

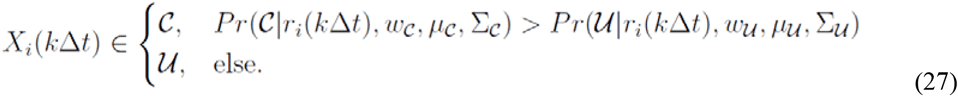

yielding a segmented trajectory where each time point is assigned as confined (C) or unconfined (U). The resulting ensembles of sub-trajectories can then be analysed further, in particular to estimate the four biophysical parameters (Eqs. 16–18, 21). In practice, if a point *X_i_*(*k*Δ*t*) ∈ C or U is isolated, e.g. C between two neighbouring U class points, then we relabel it as the class of its two immediate time neighbours (*k* − 1)Δ*t* and (*k* + 1)Δ*t*.

#### 3. Simulations to validate our 4P classification algorithm

In this section, we estimate the accuracy of the classification algorithm (subsections 2.2–2.3) using simulated (synthetic) trajectories that can switch between two states C and U. The dynamics were described by an Ornstein-Uhlenbeck equation^88^ (Eq. 28), and the generated trajectories were considered as a ground-truth ensemble, which we used to estimate the accuracy of the classification procedure developed in the previous section.

##### 3.1 Simulation of a ground-truth ensemble of synthetic trajectories

We generated *N* trajectories *X_i_*(*t*), *i* = 1,…,*n_i_*, which can switch between confined C and unconfined U states at any time *t*. The stochastic switching process is defined by:

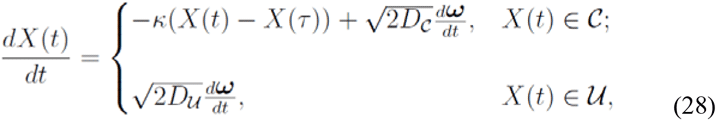

where *D*_C_, and *D*_U_ are the diffusion coefficients for the confined C and unconfined U states respectively, *d*ω/*dt* are standard three-dimensional Brownian motions, with a mean of zero and a standard deviation of one, while *κ* is the strength of a potential well that attracts the trajectory *X*(*t*) ∈ C to a fixed-point *X*(τ), which is the last position, before time *t* where the trajectory was in state U prior to switching to C:

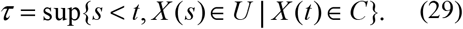

The transition probability between states (C, U) is defined by a Markov chain: The transient probability matrix *P_ij_* between state *j* at time *s* + Δ*s* and state *i* at time *s*, is given by:

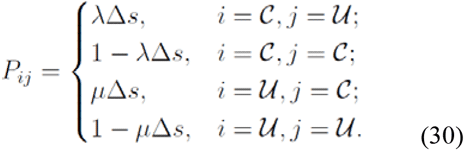

The probabilities *P*_C_(*s*) and (*P*_U_(*s*)) satisfy^93^:

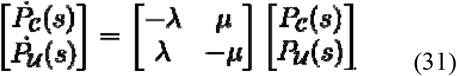

and the solution of Eq. 31 is:

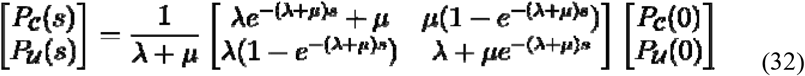

The simulation procedure was as follows: we initialized *X_i_*(0) in one of the states C or U. We used Euler’s scheme to discretize Eq. 28 at a time step Δ*t*. For each consecutive step *k*Δ*t*, *k* = 1,…*n_i_*, we determined the state of the trajectory from the probabilities *P*_C_(Δ*t*),*P*_U_(Δ*t*) (Eq. 30), using the previous state as the initial condition:

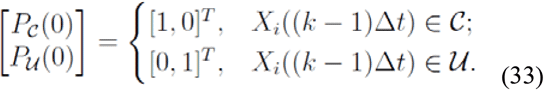

##### 3.2 Evaluating the classifier using ground-truth trajectories

To measure the accuracy of our classification algorithm (Sections 2.2–2.3), we generated *N* = 100 trajectories *Y_i_*(*k*Δ*t*), *i* = 1,…,*N*, *k* = 1,…,*n_i_* using Eq. 29, where the length of a trajectory *n_i_* was chosen randomly from a Poisson distribution with average length *n* = 100 time points, Δ*t* = 0.02*s*, *κ* = 2, different values of *D*_C_ and *D*_U_ (see below), and twenty equally spaced values of *λ*, *μ* ∈ [0,1] to obtain ground-truth states *S_i_*(*k*Δ*t*) ∈ [C,U] associated with each trajectory *Y_i_*(*k*Δ*t*). We used the four-parameter classifier (section 2.3) to classify the trajectories *Y_i_*(*k*Δ*t*) and add a label *C_i_*(*k*Δ*t*) ∈ [C, U]. To evaluate the similarity between the true class set *S_i_* and the output of the classification classes *C_i_*, we used an indicator function *δ*_*i*_(*k*Δ*t*), defined by:

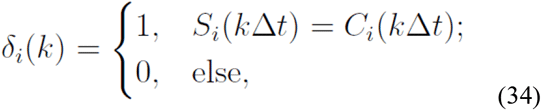

and then used Eq. 34 to define a parameter *M* that measures the accuracy of the classification along all trajectories. It was defined by:

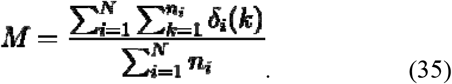

We evaluated the accuracy *M* (Eq. 35) for an ensemble of trajectories generated with twenty values of *D*_C_ ∈ [0.002, 0.1] and twenty values of *D*_U_ ∈ [0.2, 1] *μm*^2^/*s*, where we set the switching rates to be *λ* = 0.8, *μ* = 0.9): we found that the classifier achieved above 90% accuracy in the majority of the parameter space, dropping to 76% for confined and unconfined diffusion constants where we restricted *D*c ∈ [0.02, 0.1], and *D*_U_ ∈ [0.1, 0.2] [see **Figure 1b** (left), below]. In addition, to test the robustness of the classification algorithm to the switching rates, we evaluated M (Eq. 35) for twenty values of the switching rates λ, μ ∈ [0, 1] where *D*_C_ and *D*_U_ were similar (0.008 and 0.01 μm^2^/s, respectively) and obtained a very good accuracy – above 85% for all values of λ, μ [see **Figure 1b** (right), below].

**Figure 1.**
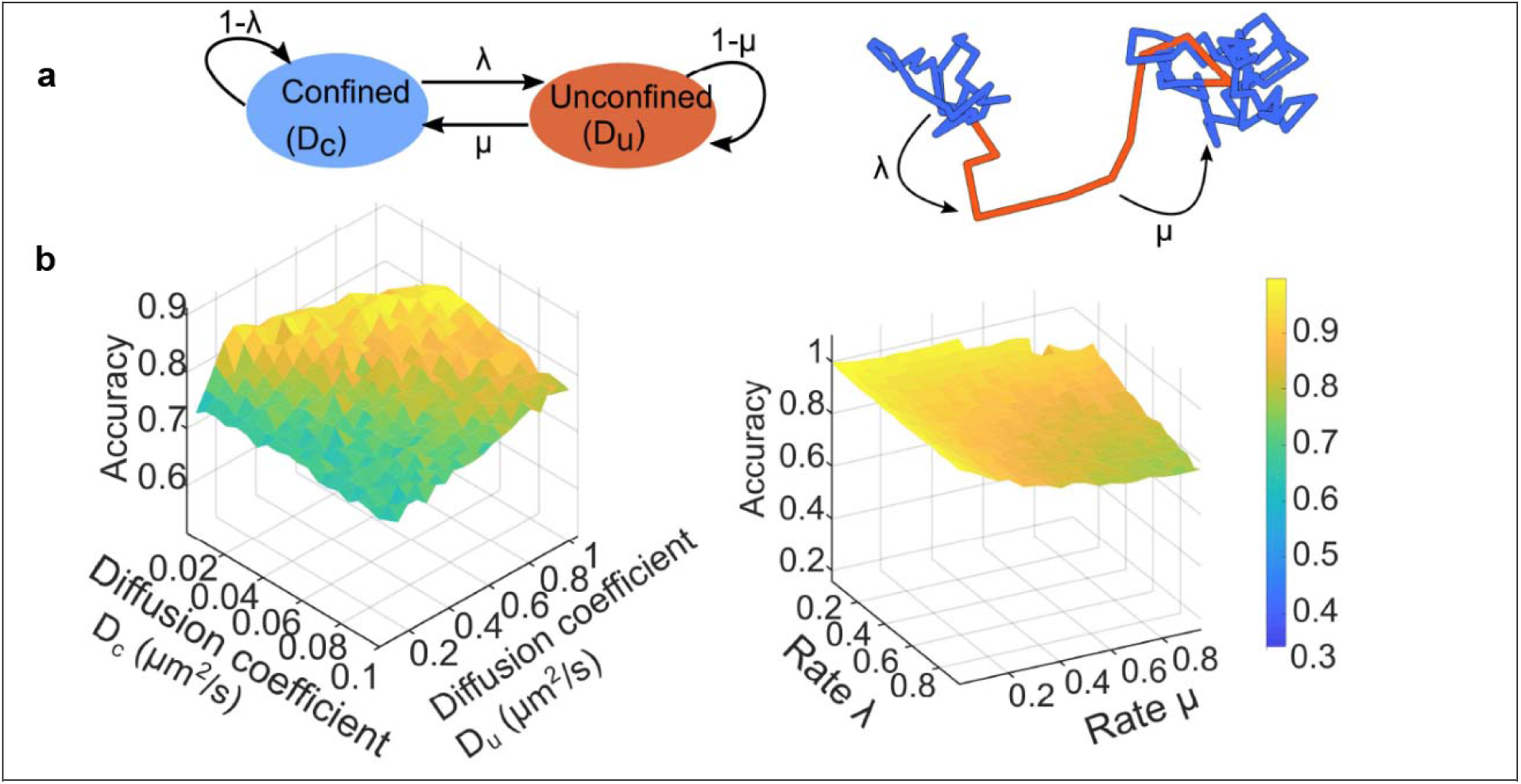
Evaluating the classifier using ground-truth trajectories with known classes (confined *C* and unconfined *U*). (a) The switching behaviour is described in Eq. 28, see above, where the class (C, U) is determined at each time point using a Markov chain (left) with switching rates *λ*, *μ* for the transitions between C and U and vice-versa, respectively (right). (b) After the ground truth set of trajectories was generated, we applied the classification procedure as described above, and computed the accuracy measure *M* (Eq. 35) defined as the fraction of correctly assigned classes by the segmentation algorithm according to the known classes in the ground-truth set. (Left) We computed *M* for trajectories generated using Eq. 29 with diffusion coefficients *D_c_* ∈ [0.002,0.1] *μm*^2^/*s* for class C and *D*_U_ ∈ [0.2,1] *μm*^2^/*s* for U. We find that *M* > 0.76 for all tested values of *D*_C_, *D*_U_, while a well separated *D_C_, D_U_* resulted in *M* > 0.9. (Right) The accuracy *M* for switching rates λ, *μ* ∈ [0,1] and *D*_C_ = 0.008, *D*_U_ = 0.01*μm*^2^/*s*, *M*, resulted in *M* > 0.85 for all tested λ, *μ*.

In summary, when tested on simulated data, the classification algorithm was able to robustly classify trajectories into confined and unconfined/sub-diffusive states.

#### 4. Estimation of association and dissociation times from trajectories

After classifying the 20 ms data trajectories into two classes C and U (see section 2.3), we estimated the association and dissociation time constants as follows. The association time *τ*_*A*_ for trajectory *X_i_* is defined as the time spent freely diffusing between confined chromatin-bound states:

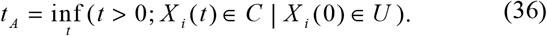

The dissociation time *τ*_*D*_ is the time spent bound to chromatin between unconfined freely diffusing states:

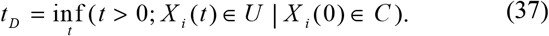

To construct the histogram of these times, we reset the origin of time to zero after each association or dissociation event. In practice, for each partition of a trajectory *X_i_* in a given ensemble, we collected the sequence 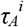 of association times as the consecutive time points for which the trajectory spends in class U between being confined, as determined by the posterior probability *P* (Eq. 26). Equivalently, the dissociation times *τ*_*D*_ are those times in which the trajectory spends in confined state C between being unconfined. We fitted the histogram of association times and dissociation times with a single exponential^94^:

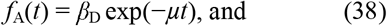

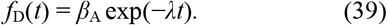

The association and dissociation times are then obtained 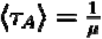 by 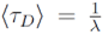 and.

#### 5. Gaussian fitting of 500 ms exposure anomalous exponents

Although the algorithm we designed segments 500 ms exposure trajectories into slow-diffusing and fast-diffusing components, the distribution of the anomalous exponents in the fast-diffusing component showed that two peaks can clearly be identified by Gaussian fitting. Indeed, when we fitted 1, 2 or 3 Gaussians to the anomalous exponent distributions for chromatin bound NuRD complex subunits such as CHD4, MBD3 and MTA2 in wild-type ES cells, we found that 2 Gaussians were the best minimal model to account for the data – having the lowest BIC value (see Eq. 13 and **Extended Data Figure 8c**). This result was supported by the initial displacement analysis that we carried out (see Section 1), which also suggested that the fast components can be decomposed into two sub-populations.

To ensure enough data had been collected to account for cell-to-cell heterogeneity, data was collated from 3 fields of view with around 6 cells in each field of view imaged, leading to a total of around 18 cells per condition. To assess the reproducibility of this result an additional 3 fields of view containing around 18 cells were collected for chromatin bound CHD4 on a different day and shown to have a similar anomalous exponent distribution (data not shown). Single molecule imaging of CHD4 in the *Mbd3-*ko was also shown to have a significantly different distribution when compared to both wild-type ES cell CHD4 datasets (p = 0.005 and p = 0.02 respectively).

#### 6. Instructions for use of the code for classification and segmentation of trajectories into two diffusion states using four parameters (4P)

To use the classification code, click “Run” on MATLAB and then select all the relevant input files to include in the classification. The input format is a .csv file with the following columns:

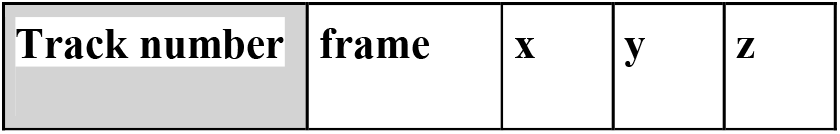

The input parameters to define are as follows:

1. Exposure time for a frame e.g. 0.02 for 20 ms or 0.5 for 500 ms

- 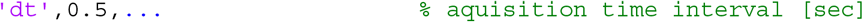
2. Options to exclude short trajectories. This parameter also specifies the minimal length that a molecule must spend confined/unconfined for that sub trajectory to contribute to association/dissociation time calculations:

- 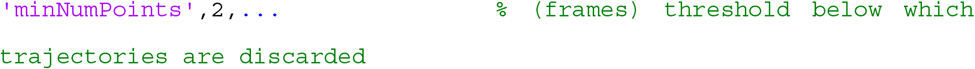
3. Option to deal with misclassification errors for accurate final parameter calculation. To increase the accuracy of the calculation, we say that the period a molecule continuously spends confined/unconfined must be at least 7 frames long, i.e. at least 7 adjacent sliding windows must be assigned to the same class:

- 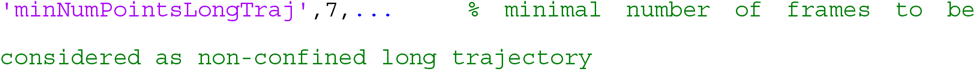
4. Number of frames to include in the sliding window either side of a point on the trajectory

- 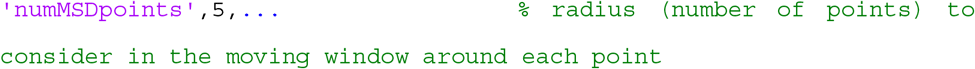
- For this study, we used optimised parameters of 5 frames before and after (11 frames in total)
- Trade-off: longer sliding windows give better estimates of the parameters, but fail to detect faster transitions
5. Option to classify by all 4 parameters or to choose a subset of them.

- 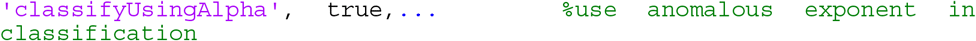
- 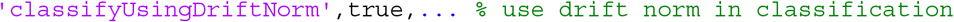
- 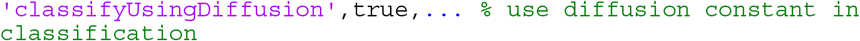
- 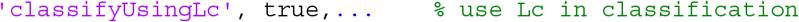
- 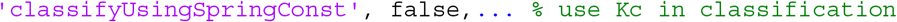

We currently do not classify by Kc, a spring constant useful for some analyses (see Shukron *et al., Trends in Genetics*, 2019), but this is included in the code in case it is of use to others.

To generate outputs, set the following export function to true:

- 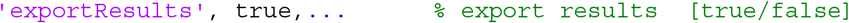

This will output the following:

1. .csv files for biophysical parameters calculated for sliding windows in each trajectory:

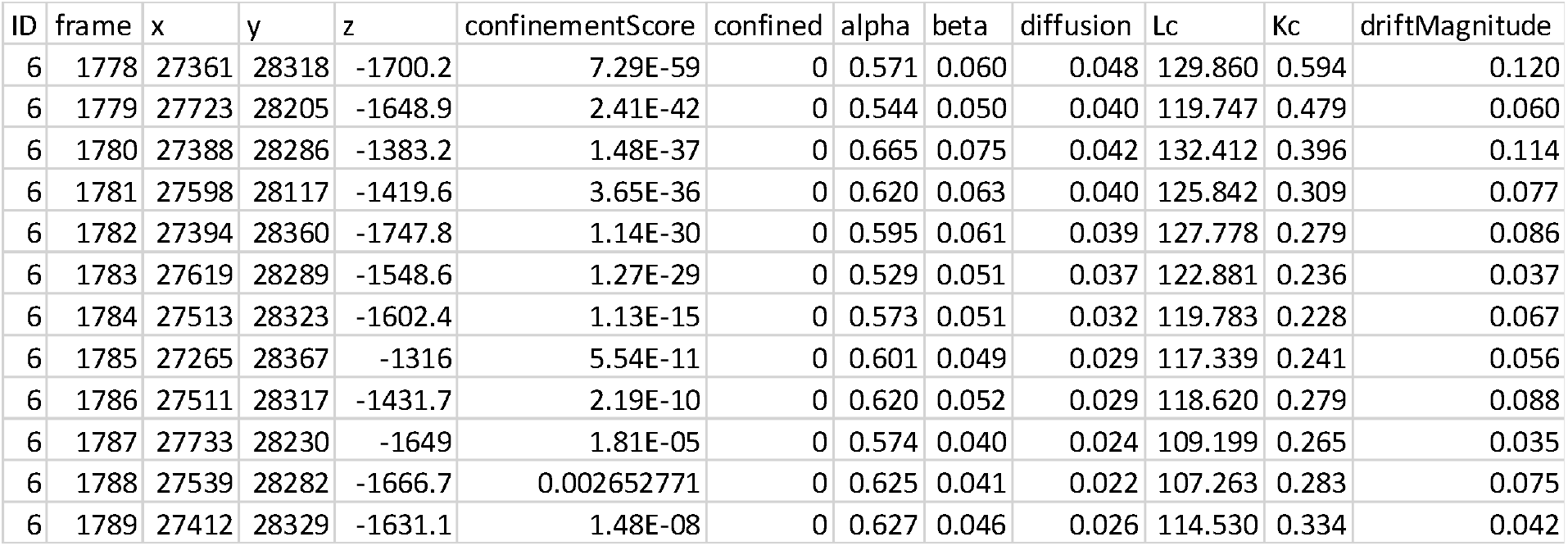

2. .csv files for each biophysical parameter calculated for periods spent confined (note that this is the average parameter for a period spent confined, so if molecules spend a longer time confined, the estimate is based on values from more sliding windows)
3. .csv files for each biophysical parameter calculated for periods spent unconfined
4. .csv file of association times (time spent unconfined between periods spent confined) – this excludes times where a molecule transitions for <10 frames as this may be a misclassification
5. .csv file for dissociation times (time spent confined between periods spent unconfined) – this excludes times where a molecule transitions for <10 frames as this may be a misclassification

Additional outputs (set to ‘true’ by default) include:

6. Visualisation function in MATLAB to see all 3D trajectories but also all confined/unconfined trajectories outputted (sub-trajectory segments are coloured blue vs orange for confined vs unconfined segments).

a. 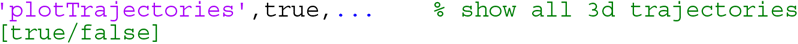
b. 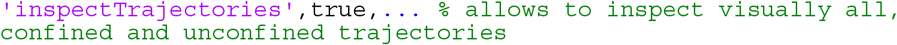
7. Options to plot the histograms of statistical parameters, association/dissociation times and trajectory duration times:

a. 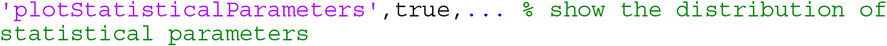
b. 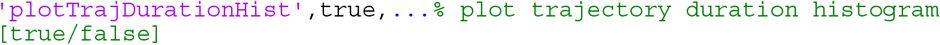
c. 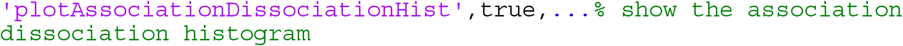
8. Option to save figures of histograms outputted in Option 7

a. 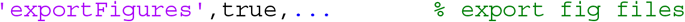
9. Export trajectories as PDB files to generate figures

a. 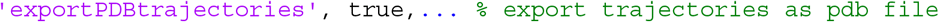
10. Changing the number of histogram fitting for association/dissociation times (default is 5)

a. 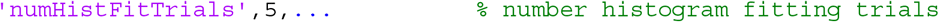
11. Option to classify by more than 2 Gaussian distributions (again not used in the paper but left as an option for other groups)

a. 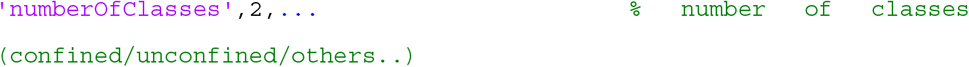

### In vitro biochemical assays of NuRD complex with and without nucleosomes

*Drosophila* PMMR and Human CHD4 were expressed in Sf21 cells and purified as described^24^. Sf21 cells expressing Human GATAD2A-MBP were resuspended in 50 mM Tris-HCl pH 7.5, 1 M NaCl, 5 mM DTT, and 1 × complete EDTA-free protease inhibitor cocktail (Roche), lysed by sonication and cleared by centrifugation at 50,000 *g* for 1 h. The supernatant was applied to amylose resin pre-equilibrated with lysis buffer and incubated for 2 h with rotation at 4 °C. The resin was washed with 20 X column volumes (CV) of lysis buffer and then eluted with 10 mM maltose in lysis buffer. Fractions containing hGATAD2A-MBP protein were concentrated and further purified by size exclusion chromatography using a Superose 6 Increase 3.2/300 column (GE Healthcare) equilibrated with 50 mM Tris-HCl pH 7.5, 150 mM NaCl, 5 % glycerol, and 1 mM DTT.

For pull-down experiments, purified protein was immobilised on MBP-Trap resin (ChromoTek) which was pre-equilibrated in pull-down buffer (50 mM HEPES pH 7.5, 300 mM NaCl, 1 mM DTT, and 5 % v/v glycerol) followed by incubation for 1 h with rotation at 4 °C. A sample of the 6% protein:bead mixture was retained as ‘Input’. The resin was washed 3 times with pull-down buffer, then a washed ‘beads’ sample was retained for analysis on a 4-12% NuPAGE gel (Invitrogen).

Electrophoretic mobility shift assays (EMSAs) were performed with n3-Widom-78bp DNA or recombinant nucleosomes made with this template^95,96^ in 10 μL of binding buffer (20 mM Hepes pH 7.5, 2 mM MgCl_2_, 5% Glycerol, and 1 mM TCEP) with varying concentrations of the indicated proteins. The reaction mixtures were incubated at 30°C for 30 min followed by centrifugation at 1000 x g. The resulting reaction mixtures were loaded onto 5% native polyacrylamide gels and run in 0.2 × TBE. Gels were stained with SYBR Gold (Invitrogen) and imaged using a Typhoon FLA 9000 (GE Healthcare).

### dCas9-GFP imaging of enhancer loci

ES cells expressing dCas9 tagged with GFP were generated as described previously^3^. Briefly, MBD3-inducible ES cells were transfected with the PB-TRE3G-dCas9-eGFP-WPRE-ubcp-rtTA-IRES-puroR vector containing a dual promoter backbone, with a TRE3G (Tet-on) promoter expressing GFP-tagged inactive dCas9 and the ubiquitin C promoter expressing the reverse tetracycline-controlled transactivator, rtTA, and a puromycin cassette via an IRES sequence. Puromycin-resistant ES cells were then selected for 7 days and doxycycline was added for 24 hours to induce expression of dCas9-GFP (through activation of the rtTA). Stable transfectants were then FACS sorted for low levels of GFP expression to select cells where only a few copies of the plasmid were integrated stably into the genome.

Prior to imaging, 1 μg/mL of doxycycline was added to ES cells for 24 hours to induce expression of low levels of dCas9-GFP. For imaging of *Tbx3* enhancer loci, three CARGO vectors in total expressing 36 gRNAs targeting the *Tbx3* enhancer were then transfected using lipofectamine 2000 (Invitrogen). The CARGO and dCas9-GFP expressing plasmids were kind gifts from the J. Wysocka lab. For imaging of *Nanog* enhancer loci, a custom designed gRNA was annealed with SygRNA® Cas9 Synthetic Modified tracrRNA (Sigma Aldrich TRACRRNA05N). The gRNA was designed such that a single gRNA sequence could be used to uniquely target a repetitive sequence close to the relevant enhancer (**Extended Data Figure 9b**). Cells were transfected with the tracr:crRNA complex using Lipofectamine 2000 (Invitrogen 11668019). In all cases, cells were transfected during passaging straight onto imaging dishes in Fluorobrite imaging medium as described above. After 24 hours, the media was replaced with fresh media and for +MBD3 samples, 4-hydrooxytamoxifen was added to a final concentration of 0.4 nM. All samples were then imaged after a further 24 hours.

2D tracking of genomic loci was carried out using oblique-angle illumination on a custom built 2D single-molecule tracking microscope as previously described^97^. Briefly, an IX73 Olympus inverted microscope was used with circularly polarized laser beams aligned and focused at the back aperture of an Olympus 1.40 NA 100× oil objective (Universal Plan Super Apochromat, 100×, NA 1.40, UPLSAPO100XO/1.4). A 561 nm laser was used as a continuous wavelength diode laser light source. Oblique-angle illumination imaging was achieved by aligning the laser off axis such that the emergent beam at the sample interface was near-collimated and incident at an angle less than the critical angle (θ_c_ ~ 67°) for a glass/water interface. This generated a ~50 μm diameter excitation footprint. The power of the collimated 488 nm beam at the back aperture of the microscope was 100 W/cm^2^. The lasers were reflected by dichroic mirrors which also separated the collected fluorescence emission from the TIR beam (Semrock, Di01-R405/488/561/635). The fluorescence emission was collected through the same objective and then further filtered using a combination of long-pass and band-pass filters (BLP01-561R and FF01-587/35). The emission signal was projected onto an EMCCD (Photometrics, Evolve 512 Delta) with an electron multiplication gain of 250 ADU/photon operating in a frame transfer mode. The instrument was automated using the open-source software micro-manager (https://www.micro-manager.org) and the data displayed using the ImageJ software^98,99^.

For image processing, PeakFit^100^ was used to localise genomic loci from the images using the filter settings: “shiftFactor”:1.0, “signalStrength”:5.0, “minPhotons”:30.0, “precisionThreshold”:40.0, “minWidthFactor”:0.5, “maxWidthFactor”:0.5, “precisionMethod”:“MORTENSEN”. Trajectories were then tracked in 2D using custom python code for connecting foci in subsequent frames if they were within 500 nm (https://github.com/wb104/trajectory-analysis). Trajectories were classified as for single molecules using a Gaussian mixture model – see **Single molecule trajectory analysis**, above.

### Enhancer-promoter DNA FISH (fluorescence in situ hybridisation)

FISH probes were prepared from mouse BAC library clones (Source Biosciences)^101^. BAC vector DNA was purified using the Qiagen Large Construct Kit. BAC DNA was labelled using Cy3 and Alexa Fluor 647 Nick Translation Labelling Kits (Jena Bioscience, PP-305S-CY3, PP-305S-AF647) and purified as previously described^17^. BAC probes were generated as follows:

| **Locus** | **BAC library code** | **Probe label** |
| --- | --- | --- |
| *Tbx3* promoter | BMQ-338M19 | Cy3 |
| *Tbx3* enhancer | BMQ-309K22 | Alexa Fluor 647 |
| *Sox2* promoter | BMQ-158G1 | Cy3, Alexa Fluor 647 |
| *Sox2* enhancer | BMQ-53J20 | Alexa Fluor 647 |
| *Bmp4* promoter | BMQ-116K2 | Cy3 |
| *Bmp4* enhancer | BMQ-249B4 | Alexa Fluor 647 |

For two-colour DNA FISH, 2 × 10^4^ cells were seeded per well on microscope slides with removable 8 well silicone chambers (Ibidi 80841). Cells were fixed in 4% formaldehyde in PBS (Pierce™ 28906, Thermo Fisher Scientific) for 10 min at RT, followed by permeabilization in 0.3% Triton X-100 (Sigma-Aldrich X100) in PBS for 15 min at RT. After 3 washes in PBS, cells were incubated in pre-warmed 2 x saline-sodium citrate (SSC) with 100 ug/mL RNAse A (Qiagen 158922) for 1 hour at 37 °C. Chambers were removed and slides washed in 2 x SSC at RT. Slides were then dehydrated using 70% ethanol, 90% ethanol and 100% ethanol for 2 mins each and left to air dry. Cells were denatured in 70% deionised formamide (Sigma-Aldrich S4117) in 2 x SSC at 80 °C for 15 minutes. Slides were then again dehydrated quickly through ice-cold 70% ethanol, 90% ethanol RT, and 100% ethanol RT for 2 mins each and again left to air dry.

For each sample slide, 150 ng of Cy3-labelled BAC probe and 150 ng of AF647-labelled BAC probe were precipitated with 5 ug of Salmon Sperm DNA (Invitrogen 15632011) using 0.1 volumes of 3M sodium acetate (pH 5.2) and 2.5 volumes of 100% ethanol. Precipitated DNA was pelleted through centrifugation at 15000 x g for 20 minutes and resuspended in 50 uL hybridization buffer (50% formamide, 10% dextran sulfate (Sigma Aldrich 42867), 0.1% SDS, 2x SSC) through incubation for 1 hour at 37 °C. Probes were denatured at 80 °C for 10 mins and then transferred to 37 °C. Sample slides were overlaid with 50 μL hybridization solution and covered with Parafilm, after which hybridization was allowed to occur at 37 °C overnight in a humidity chamber. The following day, the coverslip and hybridization solution were removed, and the slides washed four times in 2 x SSC at 40 °C for 3 mins each, then four times in 0.1 x SSC at 60 °C for 3 mins. Slides were cooled by washing in 4 x SSC at RT. After removing all the wash solution, cells were mounted in VECTASHIELD® Antifade Mounting Medium with DAPI (Vector Laboratories H-1200-10).

### Sequential IF for dCas9-GFP and DNA FISH

ES cells expressing dCas9 tagged with GFP were transfected with either CARGO plasmids or a gRNA as described above (see “*dCas9-GFP imaging…*, above)**”**. Fresh media was added after 24 hours and cells were fixed the following day in PBS containing 4% formaldehyde (Pierce™ 28908, Thermo Fisher Scientific) for 10 min at RT. Cells were permeabilised in PBS containing 0.5% Triton X-100 (Sigma Aldrich X100) for 5 min, washed three times with PBS and then treated with blocking buffer (4% bovine serum albumin (Sigma-Aldrich, A9418) in 0.1% Triton-X 100 in PBS) for 30 min. Cells were incubated with GFP-Booster Alexa Fluor^®^ 488 nanobody (ChromoTek, gb2AF488) in blocking buffer (1:1000) through incubation for 1 hour at RT. Samples were washed three times in PBS, each for 5 min.

Cells were post-fixed in PBS containing 3% formaldehyde (Pierce™, Thermo Fisher Scientific, 28908) for 10 min at RT, followed by re-permeabilization in 0.1 M HCl in 0.7% Triton X-100 in PBS for 10 min on ice. After 2 washes in 2 x SSC for 5 mins each, cells were incubated in pre-warmed 2 x SSC with 10 U/mL RNAse A (Qiagen 158922) for 1 hour at 37 °C. Slides were then equilibrated in 20% glycerol in PBS for 1 hour, followed by three consecutive freeze-thaw cycles using liquid nitrogen. After incubation for 1 hour in denaturing solution (50% formamide in 2 x SSC) at RT, slides were denatured at 70 °C for 5 mins and then washed several times in ice-cold 2 x SSC. Probes were denatured at 70 °C for 10 mins and then placed on ice to cool. Hybridisation solution was prepared with 50 ng of BAC probe and 10 μg Salmon Sperm DNA (Invitrogen 15632011) per 100 uL of hybridization buffer (50% formamide, 10% dextran sulfate, 1 mg/mL BSA, and 2 x SSC). Sample slides were overlaid with 25 uL hybridization solution per well and covered with Parafilm. After denaturation at 70 °C for 5 mins on a heat block, the slide was gradually cooled to 37 °C and hybridization allowed to occur at 37 °C overnight in a humidity chamber. The following day the coverslip and hybridization solution were removed, and the slides washed three times in 2 x SSC at 40 °C for 5 mins each, then three times in 2 x SSC at RT for 5 mins. After removing all the wash solution, cells were mounted in VECTASHIELD® Antifade Mounting Medium with DAPI (Vector Laboratories).

### IF and DNA-FISH image acquisition and analysis

Imaging was carried out using a Zeiss LSM Airy Scan 2 super-resolution microscope set for imaging of DAPI (405 nm laser, 0.8 %), Cy3 (514 nm laser, 10 %) and AF647 (639 nm laser, 90 %). Three stacks of horizontal plane images (3804 × 3804 pixels corresponding to 136.24 × 136.24 μm) with a z-step of 150 nm, were acquired for each field of view. CZI image files were then imported into IMARIS 9.6 (Bitplane) for three-dimensional modelling. Quantitative analysis of inter-probe distances within nuclei was carried out using the Surfaces and Spots modules of Imaris 9.6.

### Software and code

#### 1. Data collection

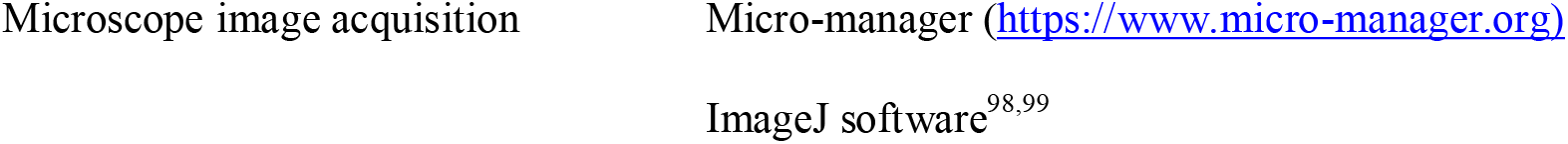

#### 2. Data analysis

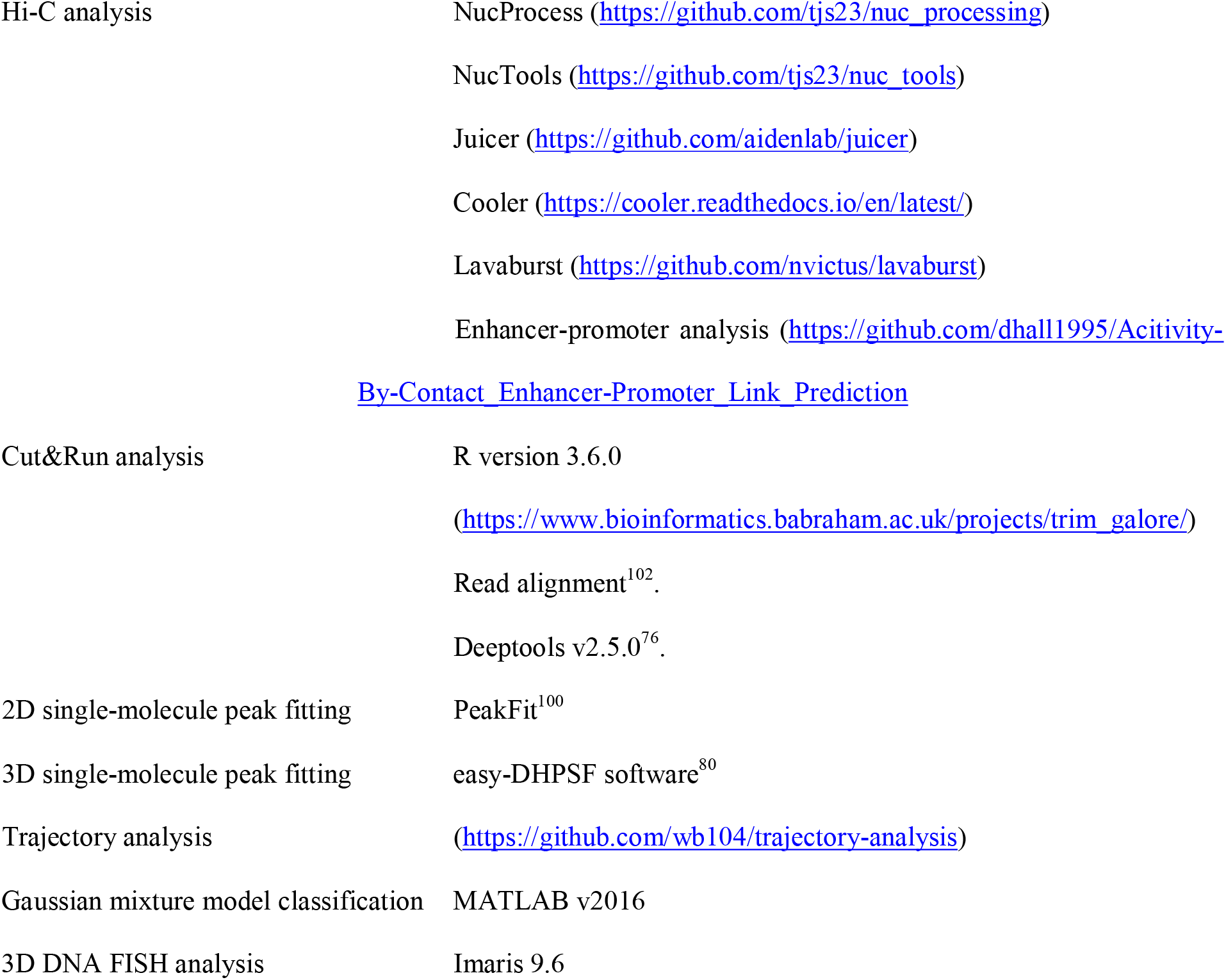

## Acknowledgements

We thank Tessa Kretschmann for preparing the figures for publication, Luke Lavis for providing the JF549 dye, Joanna Wysocka for the *Tbx3* constructs used for 2D enhancer tracking, Andy Riddell for flow cytometry and the CSCI imaging (Peter Humphreys and Darran Clements) and DNA sequencing (Maike Paramor and Vicki Murray) facilities. We thank Kathleen Bowman, Aelun Crombie and Antoine Magré for preliminary computational analysis of NuRD regulated genes, analysis of the 2D enhancer tracking experiments and KLF4 imaging, respectively. We thank the EU FP7 Integrated Project “4DCellFate” (277899), the Medical Research Council (MR/P019471/1) and the Wellcome Trust (206291/Z/17/Z) for programme funding. We also thank the MRC (MR/R009759/1 to BDH, and MR/M010082/1 to EDL), the Wellcome Trust (106115/Z/14/Z to IB and 210701/Z/18/Z to CS), and the Isaac Newton Trust (17.24(aa) to BDH) for project grant funding, and we thank the Wellcome Trust/MRC for core funding (203151/Z/16/Z) to the Cambridge Stem Cell Institute (including a starter grant to SB).

## Author contributions

SB, BDH, DHolcman and EDL designed the experiments. DL, SB, WB and TJS carried out the in-nucleus Hi-C data collection and processing. NR, WB, RR, XM and BDH carried out the Cut&Run data collection and processing. DHall, RR, XM and TJS analysed the preliminary ChIP-seq, Cut&Run and Hi-C data. LM, EB and LDC carried out preliminary ChIP-seq experiments. BDH, NR, JC, RF and SB made and characterised the cell lines used for the live-cell single-molecule imaging. ARC, AP, SFL and DK designed and built the 3D DH-PSF microscopes used for the live cell single molecule tracking experiments, and L-MN, SFL and DK designed and built the microscope for the 2D enhancer tracking. SB, DS and LHS carried out the live-cell 3D single-molecule imaging and analysed the data together with OS, PP, DK, BDH, EDL and DHolcman. SB, DS, GB, and KG carried out the live cell 2D tracking of enhancer movement. DS, SB, DL and MS carried out the immunofluorescence and DNA FISH experiments. WB developed the code to generate 3D single molecule tracks from the localisation data and dissociation time analysis, and OS, PP, AJ and DHolcman developed the code for segmentation of the 3D single-molecule trajectories. The *in vitro* biochemical experiments were carried out by WZ, JB, TD and TB, after purifying NuRD components and complexes expressed by AA, GC, IB and CS. SB, DHolcman and EDL wrote the manuscript with assistance from all the other authors.

## Data Availability

The Hi-C and Cut&Run datasets reported in this study are available from the Gene Expression Omnibus (GEO) repository under accession code GSE147789. The *Chd4*, *Klf4, Mbd3* and *Mta2* Eos-Halo targeting constructs have been deposited with Addgene.

## Code Availability

For the resubmission, we have made the data and analysis code available to the Editor and Reviewers using the following link to our group Dropbox: https://www.dropbox.com/sh/t5dm7fqb3oxdq2b/AABwXmVhOo1QL5XyPn5Fec1_a?dl=0

The password is: 5422801006. The code will be uploaded to Github upon acceptance of the paper and we will also make the source code openly available. We would also be happy to make the data available on request or to deposit it if a suitable site can be identified.

## Extended Data

**Extended Data Figure 1.**
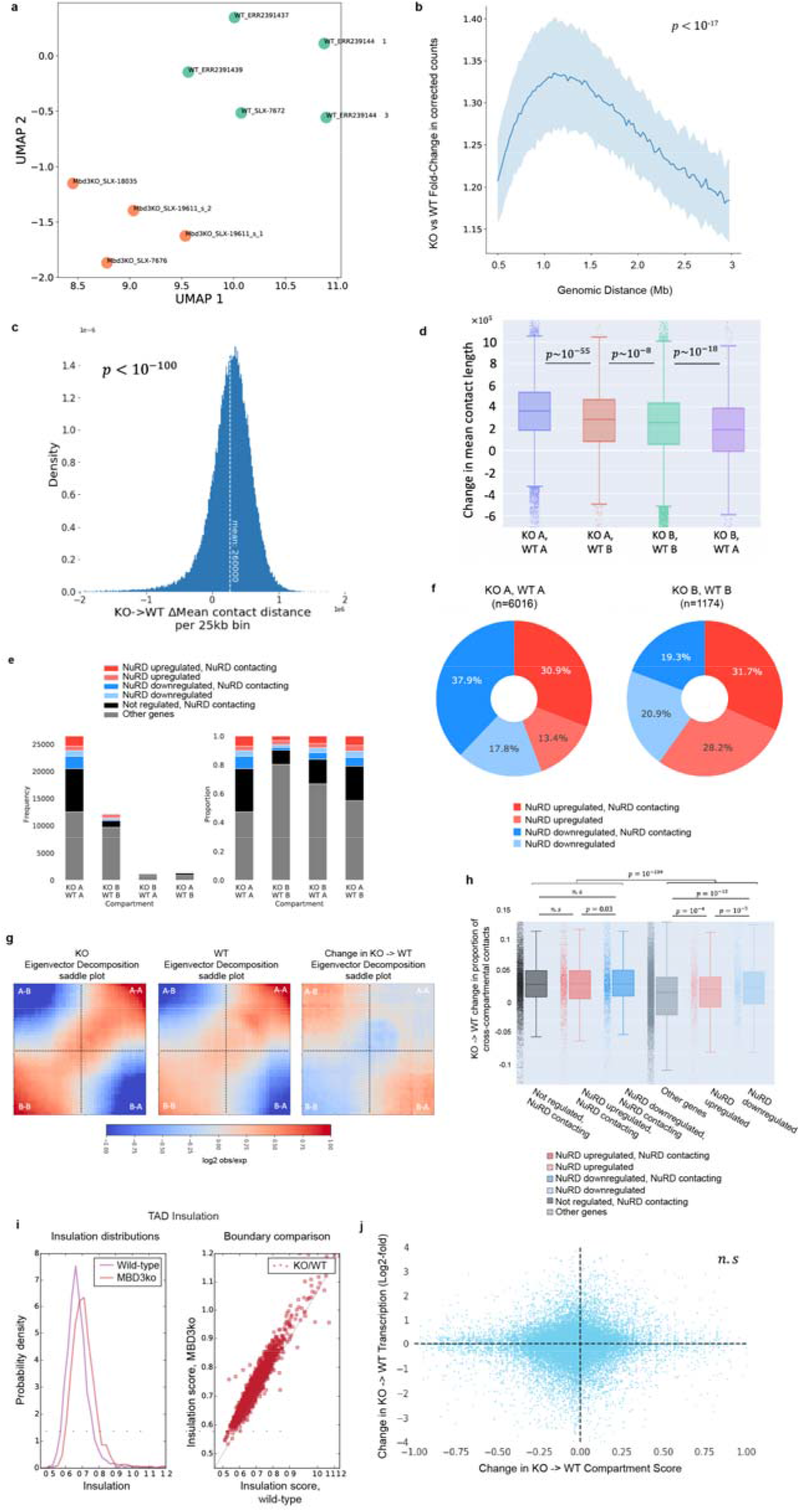
MBD3-dependent assembly of the NuRD complex increases Mb-range chromosomal interactions. (a) UMAP projection plot showing the reproducibility between our own and previously published (E-MTAB-6591) Hi-C datasets for wild-type (blue) mouse ES cells grown in 2i/LIF conditions, and comparison with those from the *Mbd3-ko* (green). (b) The fold change in numbers of contacts at a range of genomic distances shows an increase in intermediate-range (~1 Mb) contacts in the presence of the intact NuRD complex (p<10^−17^, Wilcoxon rank sum test). (c) Histogram showing the difference in mean contact distance for 25 kb genomic regions between the *Mbd3-ko* and wild-type cells. (d) Boxplot showing the changes in mean contact length for genomic regions that are in the A compartment in both wild-type and *Mbd3-ko* cells (blue), are in the B compartment in both conditions (green), that switch from A to B compartment in the presence of the intact NuRD complex (red), or that switch from B to A compartment in the presence of the intact NuRD complex (purple). (e) Bar and (f) pie charts showing the numbers of genes that remain or switch between compartments with the percentages of those genes that are up−/down-regulated. Saddle plots (g) show that there is an increase in inter-compartment contacts when going from the *Mbd3-ko* to wild-type cells, and thus A/B compartment mixing, and box plots (h) show that this is more noticeable in regions where NuRD-bound enhancers contact promoters. (i) Insulation scores derived from the contacts in the Hi-C data show a global decrease in TAD insulation in the presence of the intact NuRD complex. (j) A correlation plot shows that changes in A/B inter-compartment contacts do not correlate with changes in transcription.

**Extended Data Figure 2.**
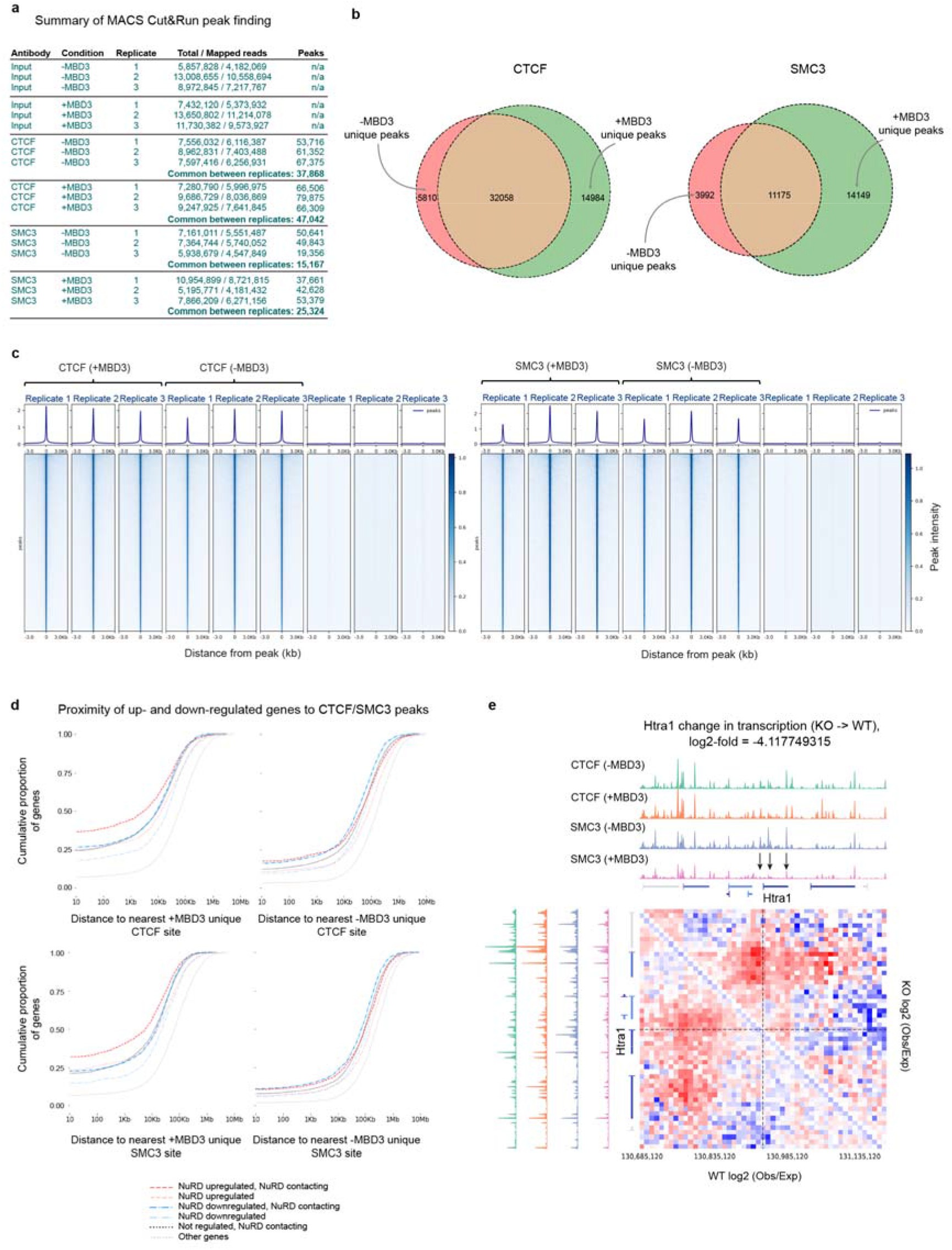
Cut&Run experiments reveal that assembly of the intact NuRD complex leads to a redistribution of both CTCF and SMC3 (a Cohesin subunit) near NuRD-regulated genes. (a) Sequencing statistics and identification of peaks that are shared in the Cut&Run experiment replicates. (b) Venn diagrams showing the overlap in CTCF and SMC3 peaks in the absence and presence of MBD3. (c) (Top) Average peak profile and (Bottom) heatmap of CTCF and SMC3 signals +/−3 kb either side of identified peaks shows no significant changes in the overall levels of CTCF and Cohesin (SMC3) in the absence and presence of intact NuRD. (d) Cumulative probability plots of the distance from different categories of promoter to the nearest CTCF or Cohesin (SMC3) binding site found uniquely in either the presence (left) or absence (right) of MBD3. These plots are compared to genes with no transcriptional change (dotted orange lines) and they all have p-values of < 1x10^−30^ (Mann-Whitney U test). (e) Comparison of the CTCF and Cohesin (SMC3) Cut&Run data, plotted together with the observed/expected contact frequencies for a given genomic distance around the *Htra1* gene. Positions where Cohesin is lost within the body of the *Htra1* gene and upstream of its promoter are indicated with black arrows. The contact maps for the *Mbd3-ko* and wild-type cells are shown above/below the diagonal, respectively, and they are coloured according to their log-fold-change in contact frequency (red = increased, blue = decreased, intensity = absolute log fold change). The genes are coloured according to their log-fold-change in levels of expression (red = upregulated, blue = downregulated, intensity = absolute log fold change) and the black dotted lines mark the position of the *Htra1* promoter.

**Extended Data Figure 3.**
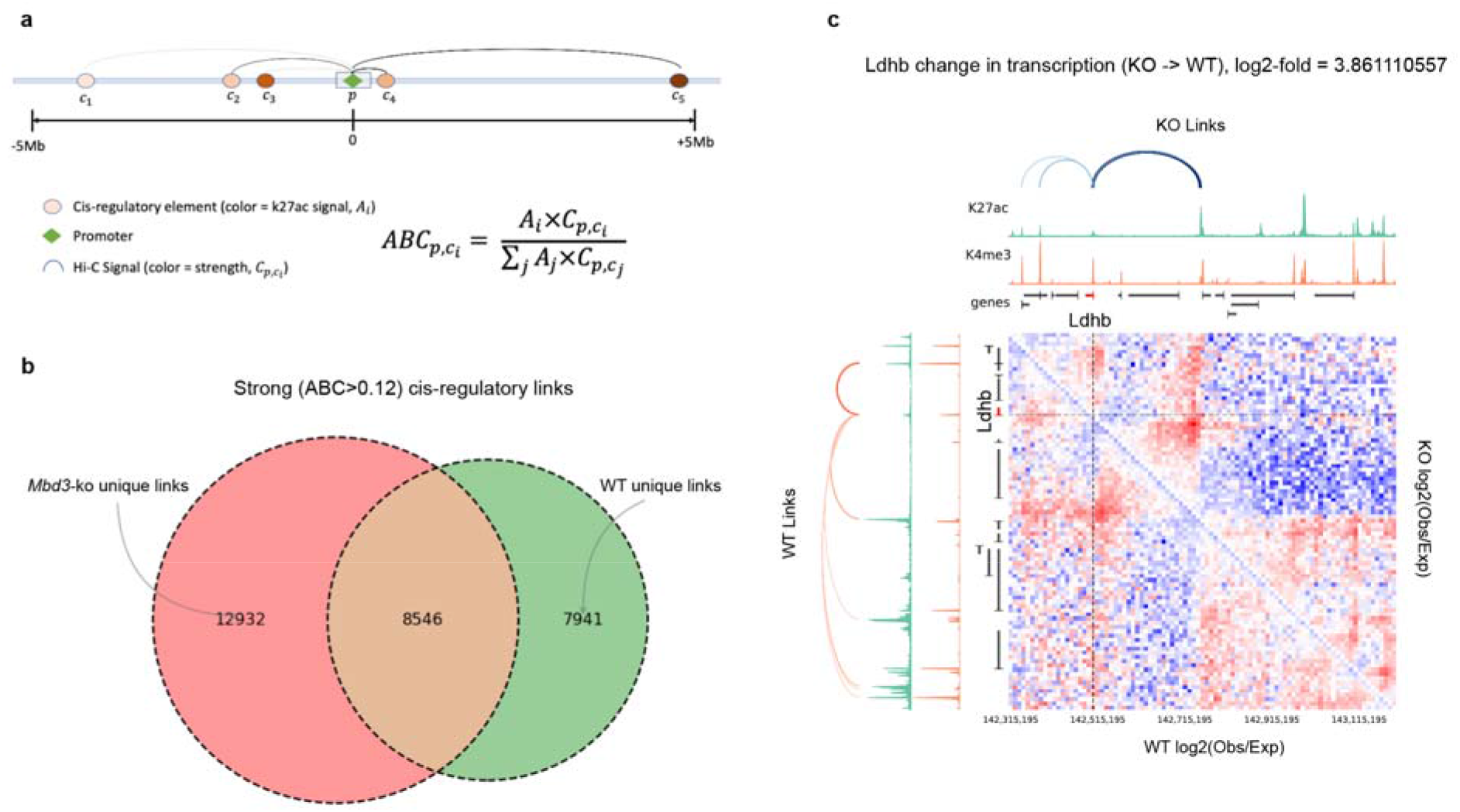
Activity-by-contact model analysis of enhancer-promoter interaction strength in *Mbd3* knockout and wild-type ES cells. (a) Schematic of the activity-by-contact (ABC) analysis illustrating how the enhancer-promoter (E-P) interaction strength (ABC score) is calculated from the H3K4me3 signal at promoters and H3K27ac signal at nearby enhancers (defined using our previously published ChIP-seq data in wild-type and *Mbd3*-depleted ES cells)^18^ as well as the Hi-C signal strength. (b) Venn diagram showing the overlap between strong enhancer-promoter contacts identified in *Mbd3* knockout and wild-type ES cells. (c) Contact maps for a region around the *Ldhb* gene with changes in enhancer-promoter interactions and loops/TADs indicated. The maps for the *Mbd3-ko* and wild-type cells are shown above/below the diagonal, respectively, and they are coloured according to their log-fold-change in contact frequency relative to that expected theoretically at that particular distance (red = increased, blue = decreased, intensity = absolute log fold change). The black dotted lines mark the position of the *Ldhb* promoter.

**Extended Data Figure 4.**
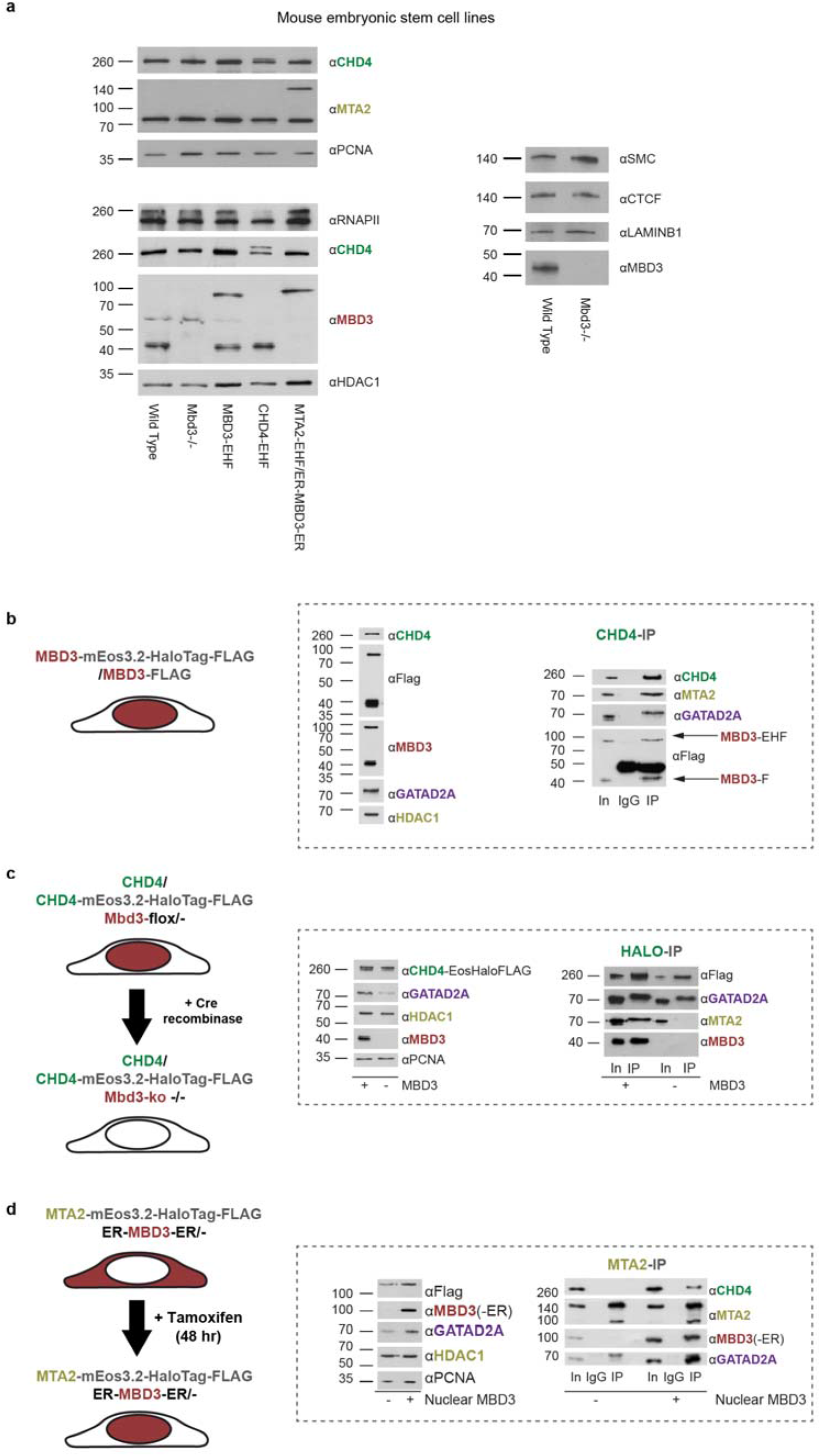
Mouse embryonic stem cell lines expressing mEos3.2-HaloTag-FLAG tagged NuRD complex subunits. (a) Western blot comparison of expression of (Left) NuRD components, and (Right) CTCF and Cohesin in the cell lines used. Detailed schematic of the (b) *Mbd3*, (c) *Chd4* and (d) *Mta2* cell lines generated. CHD4 was tagged as previously described^24^. MTA2 was tagged in ES cells expressing the ER-MBD3-ER (estrogen receptor-MBD3-estrogen receptor) fusion protein so that nuclear localisation of MBD3 is tamoxifen-inducible^18^. (Left) Expression of NuRD complex subunits was confirmed by western blot. Note that the stability of MTA2 and GATAD2A are both dependent upon MBD3, but that of CHD4 is not^68^. (Right) Immunoprecipitation of either CHD4 or MTA2 confirms that the Eos-Halo-FLAG tags do not prevent association with other NuRD components, and that NuRD complex integrity is dependent upon the presence of MBD3.

**Extended Data Figure 5.**
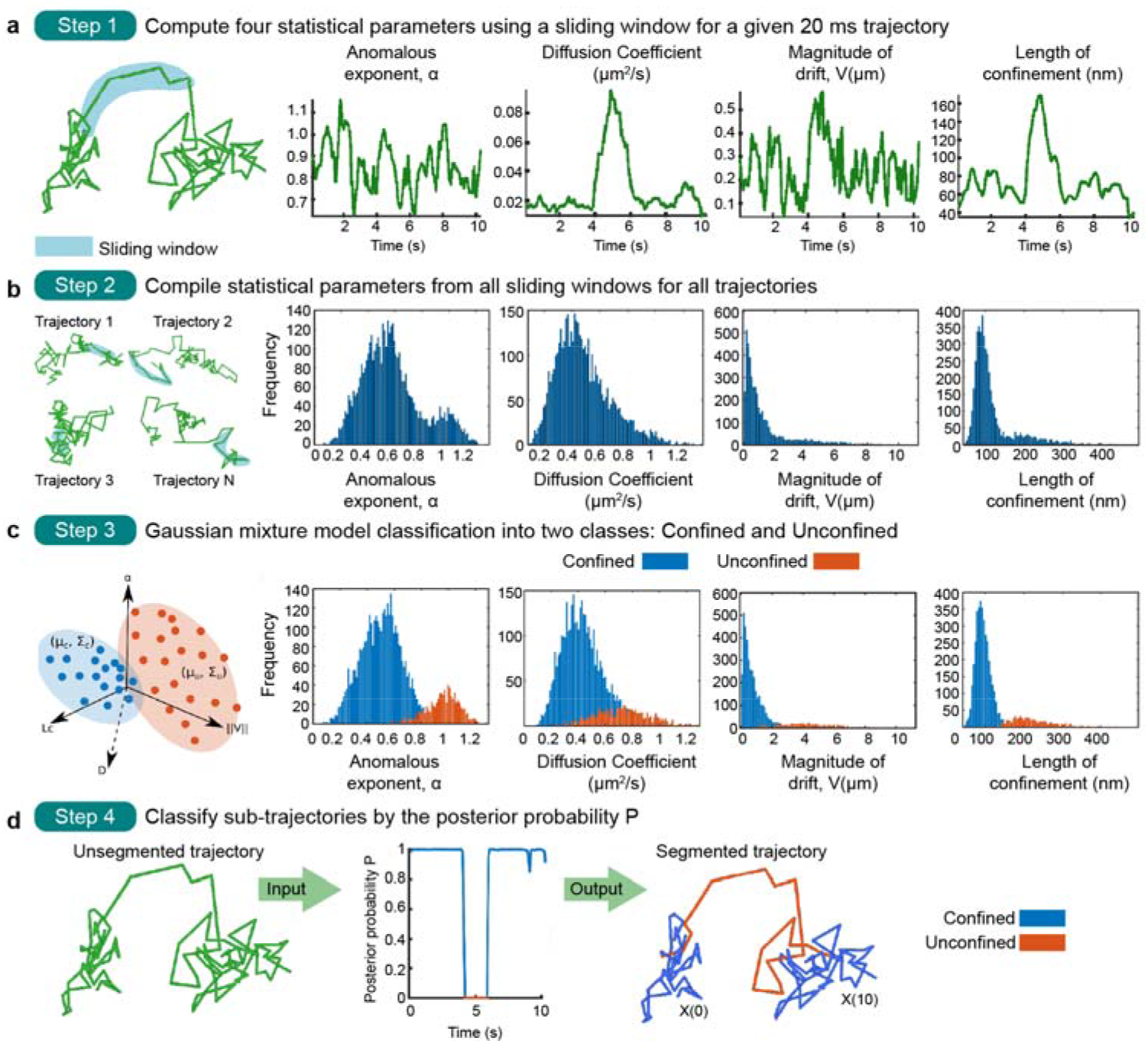
Segmentation of single-molecule trajectories of CHD4-HaloTag-JF_549._ (a) (Left) A single molecule trajectory, where we show an example sliding window (blue). (Right) Four biophysical parameters are extracted from sliding windows within this trajectory: the anomalous exponent α, the effective diffusion coefficient D, the norm∥V∥ of the mean velocity, and the length of confinement Lc, were all estimated from a sliding window of size w=10 Delta t (see Online Methods). (b) (Left) Several trajectories with example sliding windows (blue). (Right) Histograms of the biophysical parameters extracted in (a) from individual sub-trajectories are computed over the ensemble of all the recorded trajectories. (c) (Left) Scheme of the Gaussian mixture model (GMM) used to separate the histograms of the four parameters (four-dimensional feature space) into confined (C) and unconfined (U) populations (for 20 ms trajectories). The mean vectors μC, μU and covariance matrices ΣC, ΣU are estimated from the maximum likelihood. (Right) Separation of histograms in (b) into confined (blue) and unconfined (orange) populations. (d) Classification procedure applied to each trajectory, resulting in confined (blue) and unconfined (orange) sub-trajectories. The posterior probability P of the GMM (panel c) is computed on the four parameters for each point [Xi(kΔt) ∈ C with P(kΔt) > 1 − P(kΔt) (blue); otherwise X(kΔt) ∈ U (orange)] (see Online Methods). The result is a segmented trajectory where each time point is assigned as confined (C) or unconfined (U).

**Extended Data Figure 6.**
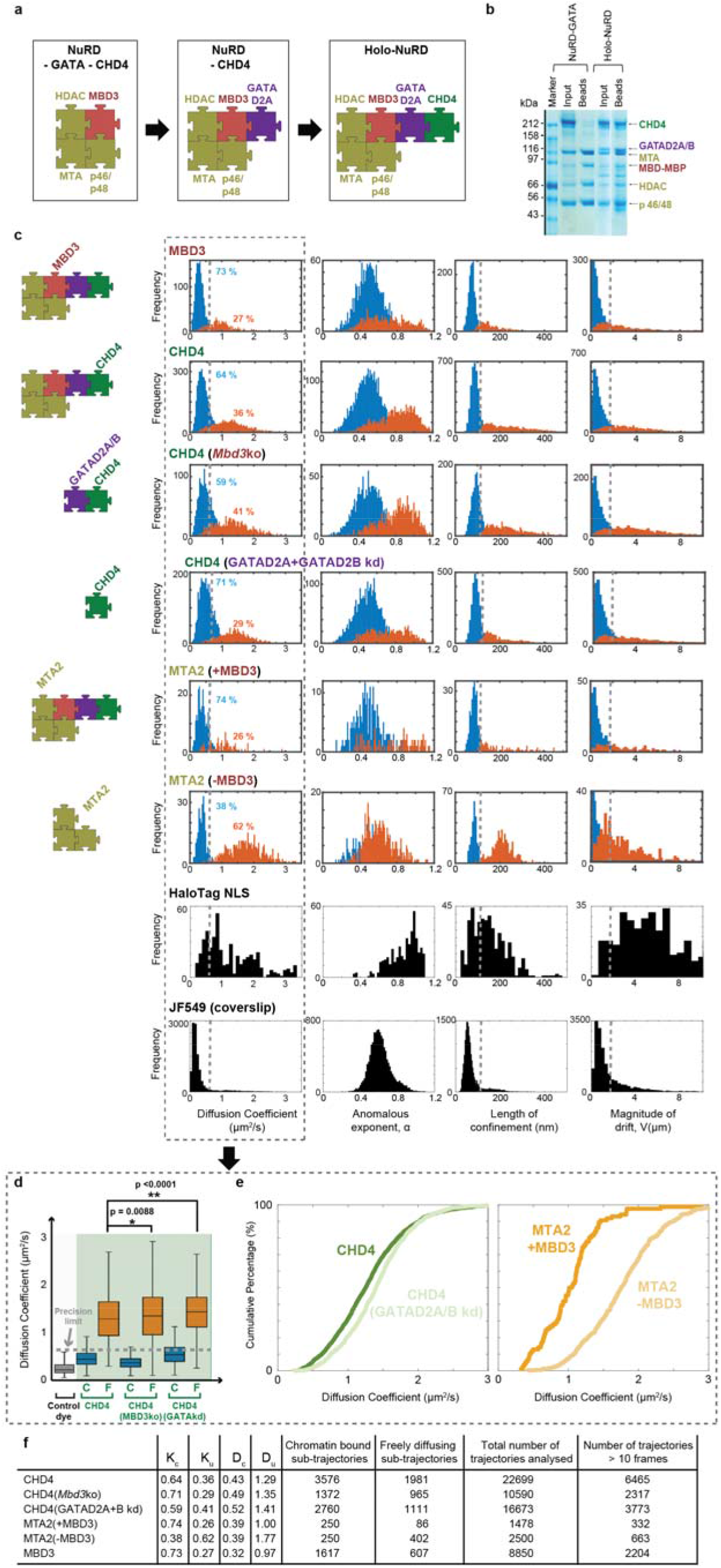
*In vitro* and live cell single-molecule imaging experiments delineate holo-NuRD complex assembly. (a) Schematic of holo-NuRD complex assembly with GATAD2A linking MBD3 to the CHD4 remodeller. (b) Pull-down experiments of MBP-tagged MBD with and without GATAD2A confirm that GATAD2A is required for CHD4 to interact with the deacetylase sub-complex. (c) Distribution of the four biophysical parameters described in **Figure 2** for 20ms exposure tracking of MBD3 and CHD4 in wild-type ES cells, as well as CHD4 in the absence of either MBD3 or GATAD2A/B. The data for MTA2 in the presence and absence of nuclear localised MBD3 are also shown. The grey dotted lines indicate the upper bound of the precision limit calculated at the 95 % confidence interval for an immobilised JF_549_ dye control sample. HaloTag with a nuclear localisation sequence (HaloTag-NLS) is also shown as a control for a (mostly) freely diffusing molecule^57,58^. (d) Box plot of apparent diffusion coefficients extracted from chromatin bound (C) and freely diffusing (F) CHD4 molecules in wild-type, *Mbd3* knockout and GATAD2A/B knock-down ES cells (*p < 0.01, **p < 0.001, Kolmogorov-Smirnov test). The grey dotted line indicates the upper bound of the precision limit calculated at the 95 % confidence interval for an immobilised JF_549_ dye control sample. (e) Cumulative distribution functions showing a higher diffusion coefficient for freely diffusing unconfined CHD4 upon removal of GATAD2A/B, and for freely diffusing MTA2 molecules upon removal of MBD3 from the nucleus. (f) Table showing the proportions (K) and estimated values of the apparent diffusion coefficients (D), as well as the number of chromatin bound and freely diffusing sub-trajectories obtained from the total number of trajectories analysed. (NB – many trajectories were discarded as they were either too short for analysis or because they had a low probability of being classified as confined or unconfined.)

**Extended Data Figure 7.**
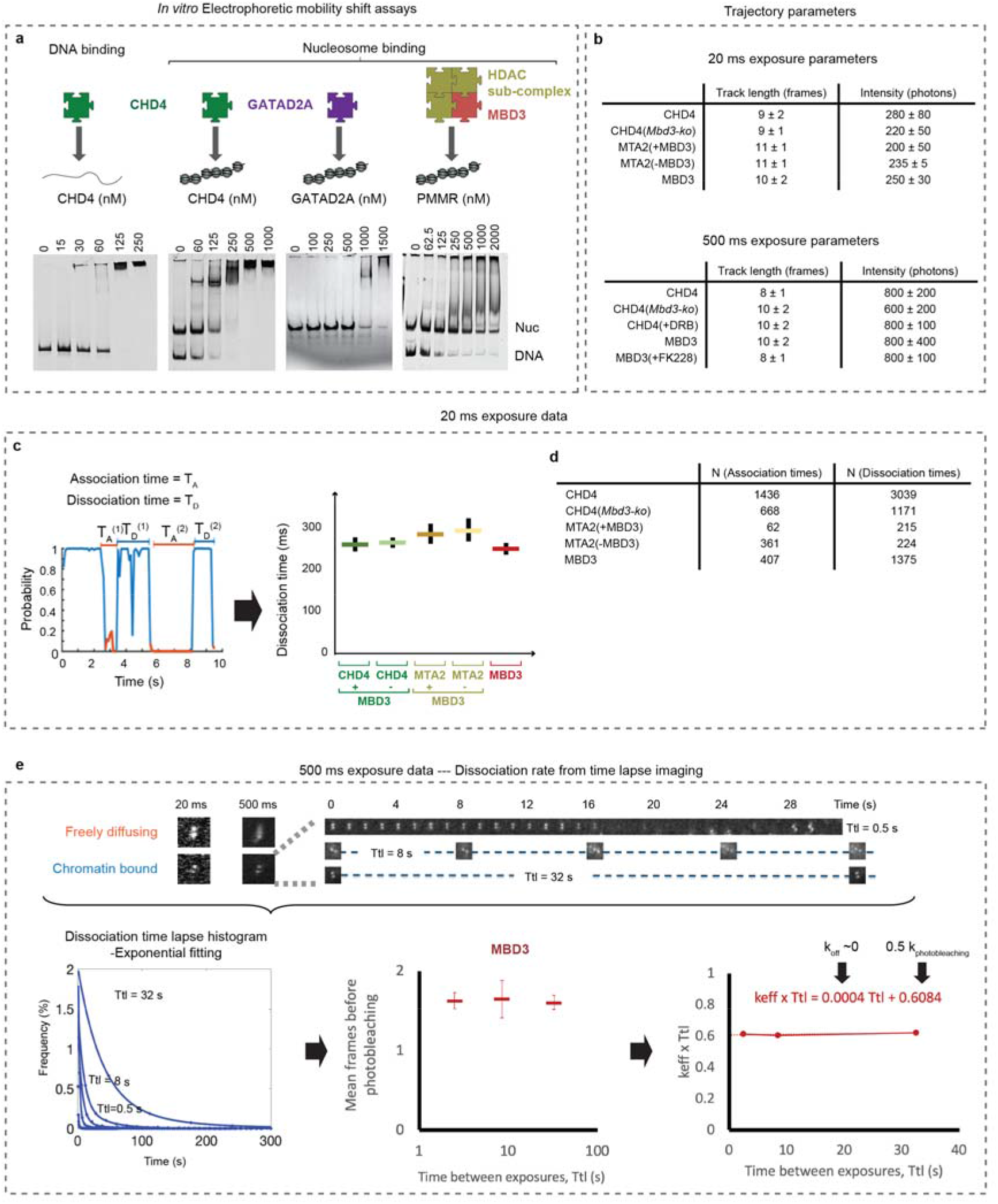
The NuRD complex interacts with both DNA and nucleosomes, but only forms tight interactions through the CHD4 remodeller. (a) *In vitro* electrophoretic mobility shift assays confirm that CHD4 binds to both DNA and nucleosome core particles (NCPs) to form large complexes that only just enter the gel. GATAD2A alone shows low affinity binding to NCPs whilst the deacetylase complex interacts with DNA, but does not bind stably to NCPs. (b) Table showing the mean track length in frames, and mean photons detected per image frame, in 20 ms (Top) and 500 ms (Bottom) exposure trajectories. (c) (Left) Confinement probability allows collection of the association T_A_ or dissociation T_D_ times – defined respectively as the time a trajectory spends between periods of confined or unconfined motion. (Right) Dissociation times calculated using transitioning trajectories as periods of confined motion between two periods of unconfined motion (see also **Figure 3**). Error bars show 95 % confidence intervals. (d) Table with the number of single molecule tracks that were used to determine the association and dissociation times. (e) (Top) Example images demonstrating how long 500 ms exposures motion blur freely diffusing molecules, but allow detection and tracking of those that are chromatin bound. Images of single chromatin bound CHD4 molecules during time-lapse imaging with various dark times. (Bottom) Exponential fitting of time-lapse residence time histograms can be used to extract the photobleaching rate k_b_ and the effective dissociation rate k_eff_. However, examination of the mean number of frames before photobleaching for MBD3, where the time between exposures is varied, shows that the results are completely dominated by photobleaching.

**Extended Data Figure 8.**
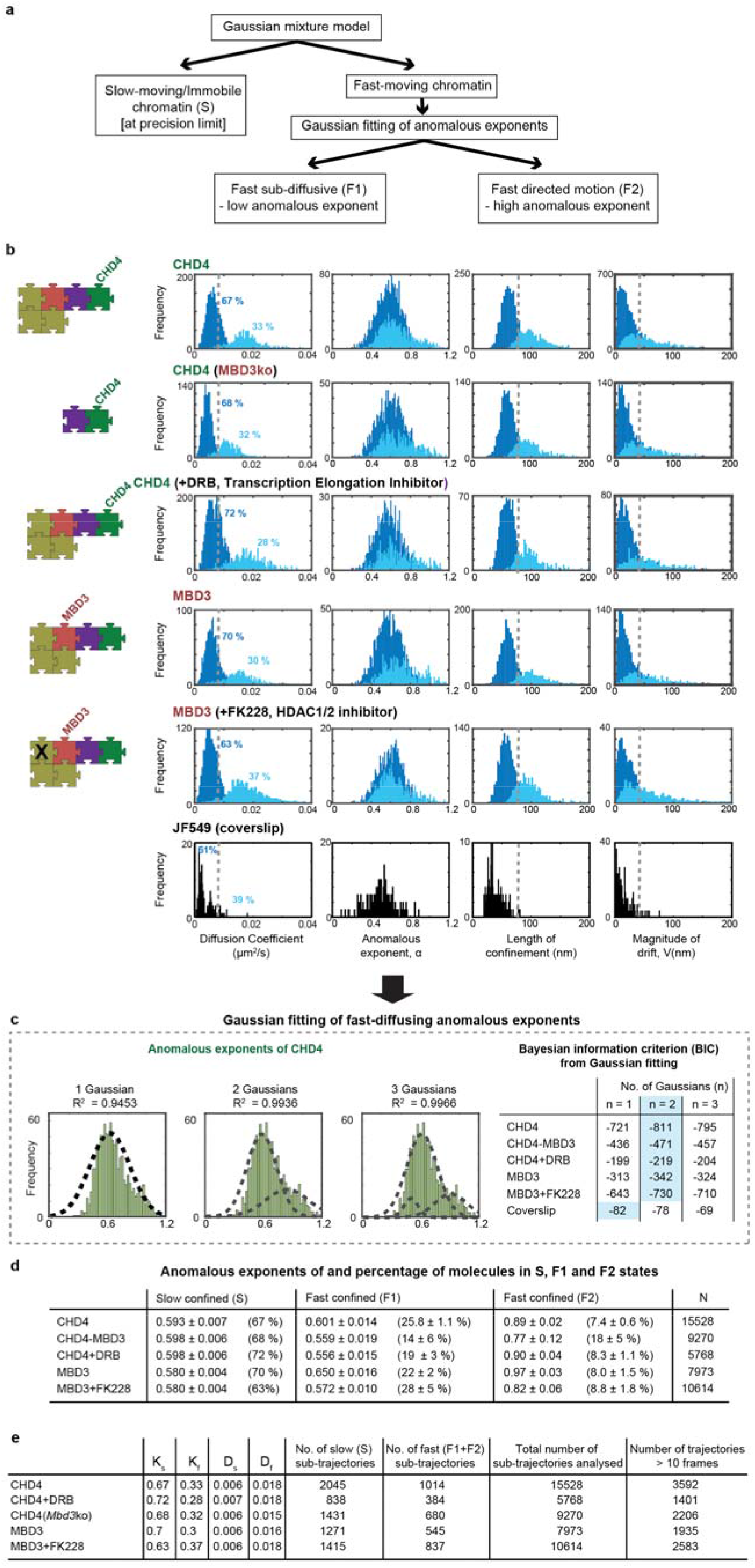
Live cell single-molecule imaging experiments to study the chromatin-bound NuRD complex. (a) Schematic of the approach used for analysing the 500 ms exposure trajectories of chromatin bound NuRD complex subunits. First, a Gaussian mixture model is used to distinguish slow-moving/immobile and fast-moving chromatin bound sub-trajectories based on the 4 biophysical parameters (see also **Figure 2** and **Extended Data Figure 5**). Then, Gaussian fitting of the anomalous exponent distributions of the fast-moving chromatin bound sub-trajectories revealed two fast-moving populations (F1 and F2) characterised by distinct anomalous exponents. (b) Distribution of the four biophysical parameters for 500 ms exposure tracking of: (i) chromatin bound CHD4 in wild-type ES cells, in the absence of MBD3, and in the presence of DRB (an inhibitor of transcriptional elongation); (ii) chromatin bound MBD3 in wild-type ES cells, and in the presence of the HDAC1/2-specific inhibitor FK228; and (iii) JF549 dye bound to the coverslip. The grey dotted lines indicate the upper bound of the precision limit calculated at the 95 % confidence interval for the immobilised JF_549_ dye control sample. (c) (Left) Fitting of 1, 2 or 3 Gaussians to the anomalous exponent distributions for fast moving chromatin bound CHD4 in wild-type ES cells – the R^2^ values above indicate the goodness of fit. (Right) The Bayesian information criterion (BIC) was calculated for all the datasets shown in (b) to determine that 2 Gaussians are the best minimal model to account for the data – i.e. that has the lowest BIC value (light blue box). (d) Table summarising the changes in anomalous exponent of the slow and fast chromatin bound NuRD complex subunits in the presence and absence of MBD3, or in the presence of specific inhibitors. Errors given are for 95 % confidence intervals. (e) Table showing the proportions (K) and estimated values of the apparent diffusion coefficients (D), as well as the number of slow (S) and fast (F1+F2) diffusing sub-trajectories obtained from the total number of trajectories analysed. (NB – many trajectories were discarded as they were either too short for analysis or because they had a low probability of being classified as slow or fast.)

**Extended Data Figure 9.**
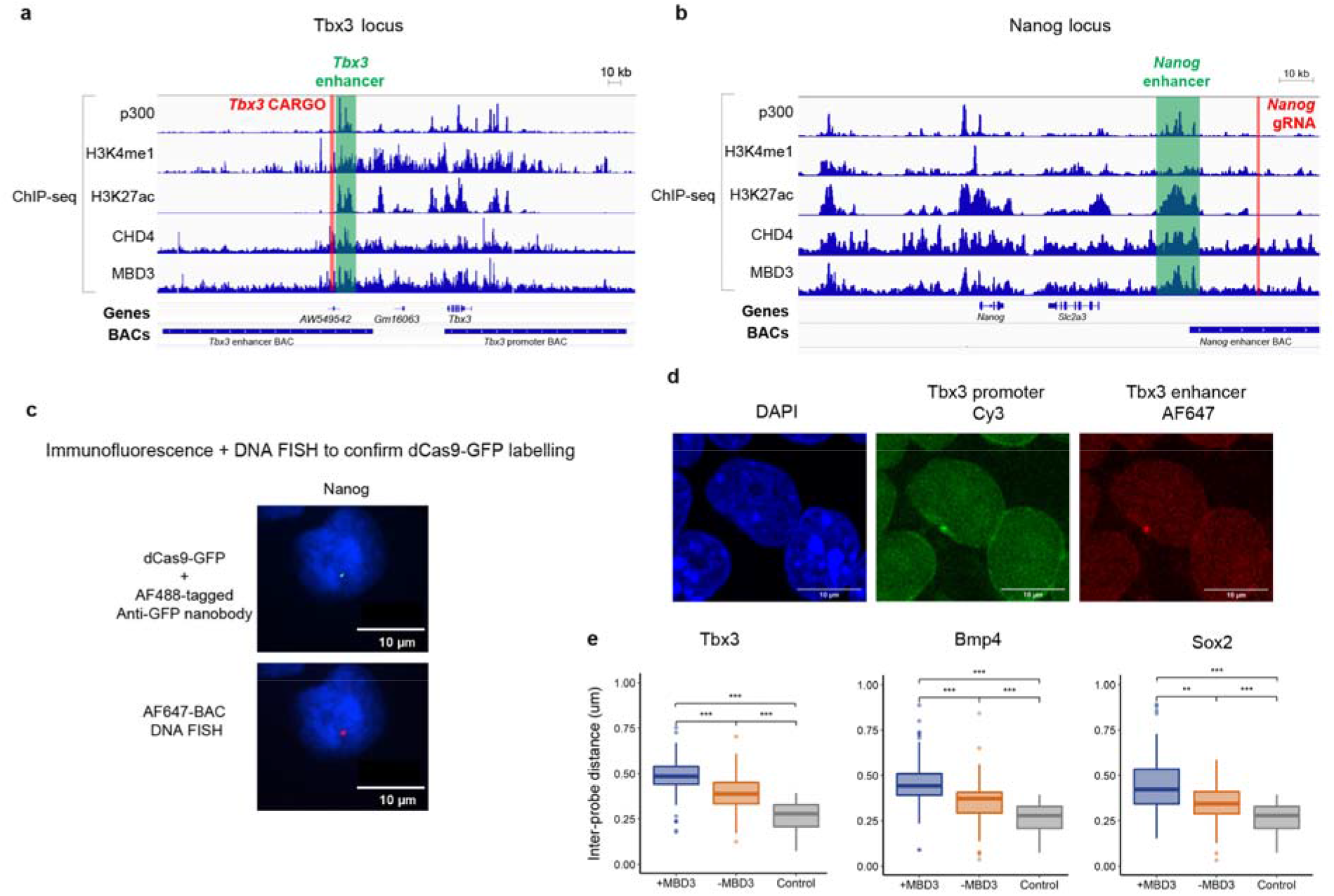
DNA FISH validation to confirm NuRD-dependent changes in enhancer-promoter compaction. Genomic locations of the (a) *Tbx3* and (b) *Nanog* genes annotated with the locations to which dCas9-GFP was targeted using either CARGO vectors^3^ or a single gRNA that targets nearby genomic repeats (red lines). Locations of bacterial artificial chromosome (BAC) DNA FISH probes for the *Tbx3* and *Nanog* enhancers are also indicated as are the targeted enhancers themselves (green). The corresponding ChIP-seq profiles^17^ indicate the binding of the NuRD complex subunits CHD4 and MBD3 as well as the location of active enhancers (determined from the ChIP-seq profiles for H3K27ac, H3K4me1 and p300). (c) Representative confocal images showing co-localisation of dCas9-GFP (labelled using AF488-tagged anti-GFP nanobody) and AF647-BAC DNA FISH probes targeting the *Nanog* enhancer. (d) Representative confocal images of Cy3-labelled BAC DNA FISH probes targeting the *Tbx3* promoter and AF647-labelled BAC DNA FISH probes targeting the *Tbx3* enhancer. (e) Boxplots showing the enhancer-promoter distances in MBD3-inducible ESCs with and without tamoxifen: +MBD3 (blue) and -MBD3 ESCs (orange) respectively. There is a significant increase in enhancer-promoter distance in the presence of intact NuRD for *Tbx3* (+MBD3, n = 70; -MBD3, n = 101), *Bmp4* (+MBD3, n = 172; -MBD3, n = 71) and *Sox2* (+MBD3, n = 42; -MBD3, n = 50) (**p<0.01, ***p < 10^−5^, Kolmogorov-Smirnov test). To estimate the precision limit of the experiment, control samples were generated in which the distance was measured for the *Sox2* enhancer labelled with both Cy3 and AF647 (grey, n = 32).

**Extended Data Figure 10.**
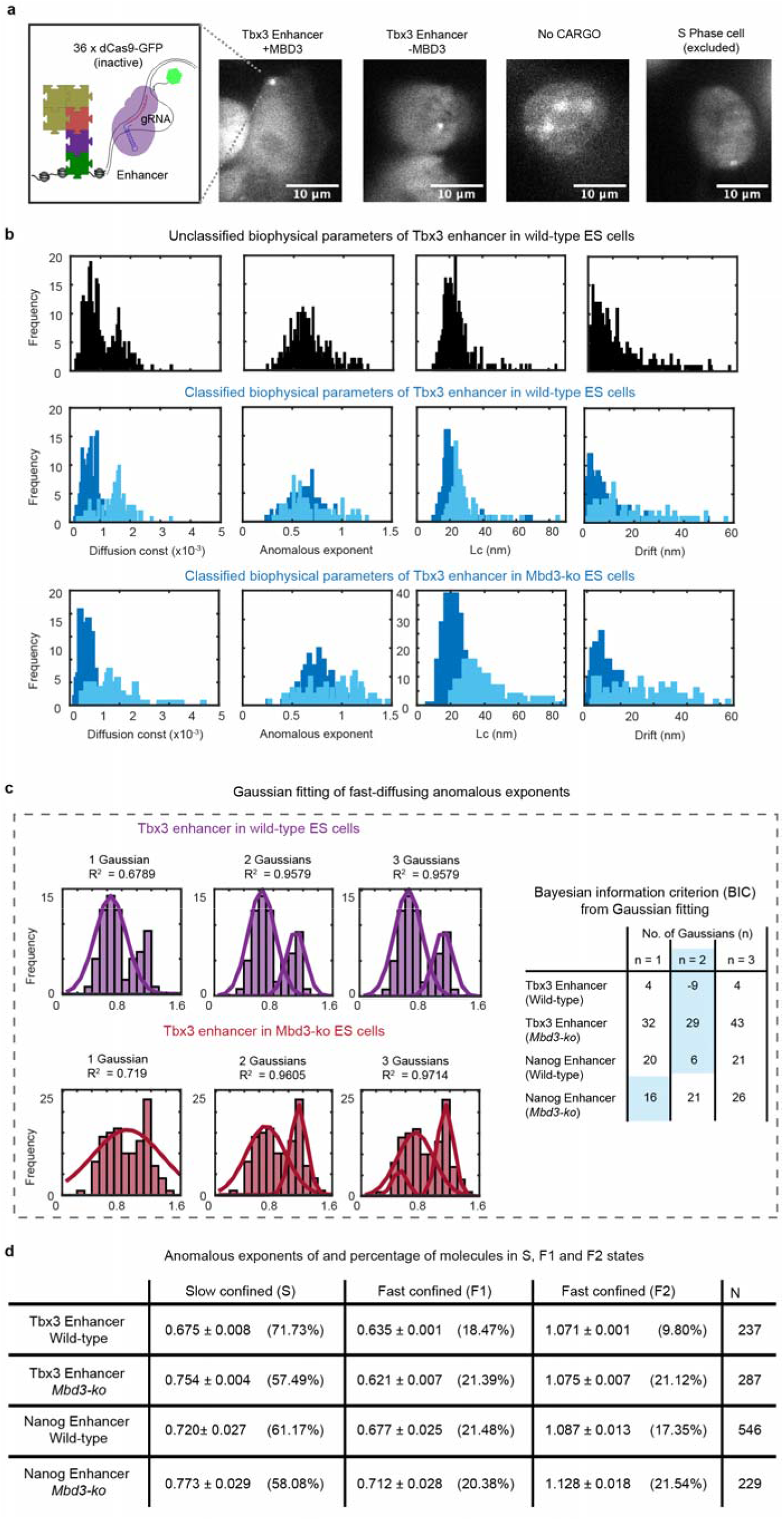
2D dCas9-GFP tracking of active enhancers. (a) Representative images of 36 gRNAs targeted to the *Tbx3* enhancer in the presence or absence of MBD3 with a negative control expressing no gRNAs. (Right) An example of a cell with a doublet indicating that it is in S phase (that was excluded from the analysis) is also shown. (b) Distribution of the four biophysical parameters extracted from sliding windows within the 2D single-molecule trajectories of dCas9-GFP bound at the *Tbx3* enhancer in wild-type ES cells imaged using 500 ms exposures – (Top) before and (Middle) after classification based on the anomalous exponent α, the apparent diffusion coefficient D, the length of confinement Lc, and the drift magnitude, norm∥V∥ of the mean velocity. (Bottom) Distribution of the four biophysical parameters after classification for the *Tbx3* enhancer in *Mbd3-ko* cells. (c) (Left) Fitting of 1, 2 or 3 Gaussians to the fast-moving anomalous exponent distributions of the *Tbx3* enhancer tracked in either wild-type ES (Top) or *Mbd3-ko* cells (Bottom) – the R^2^ values above indicate the goodness of fit. (Right) The Bayesian information criterion (BIC) was calculated for all the datasets to determine which number of Gaussians best modelled the data – that with the lowest BIC value (light blue box). (d) Table showing the Gaussian fitted anomalous exponent values for slow- and the fast-moving chromatin bound dCas9-GFP at both enhancers tracked in the presence and absence of MBD3.

## Notes

### Competing Interest Statement

The authors have declared no competing interest.

### Summary of Updates

Updated analysis and added data required by reviewers

